# PARP1 and PARP2 are dispensable for DNA repair by microhomology-mediated end-joining during mitosis

**DOI:** 10.1101/2025.06.09.658719

**Authors:** Raquel Ortega, Erin Taylor, Sophie M. Whitehead, Thomas Danhorn, Benjamin G. Bitler, Nausica Arnoult

**Affiliations:** Department of Molecular, Cellular, and Developmental Biology, University of Colorado Boulder, Boulder, Colorado, 80309, USA; Division of Reproductive Sciences, Department of Obstetrics and Gynecology, University of Colorado Denver, Anschutz Medical Campus, Aurora, Colorado, 80045, USA; University of Colorado Cancer Center, University of Colorado Anschutz Medical Campus, Aurora, Colorado, 80045, USA; Department of Biomedical Informatics, University of Colorado Anschutz Medical Campus, Aurora, Colorado, 80045, USA

## Abstract

Poly ADP-ribose polymerase (PARP) inhibitors are standard of care treatment for cancers with homologous-recombination deficiencies. Yet, as tumors develop resistance, complementary strategies are emerging, including targeting microhomology-mediated end joining (MMEJ). Given that PARP1 is widely described as a key promoter of MMEJ, one would expect PARP inhibition to suppress MMEJ—potentially making MMEJ inhibition redundant. MMEJ was originally described as a backup pathway, with seminal work linking PARP1 to MMEJ conducted in NHEJ-deficient cells, where MMEJ can be reactivated in G1. However, we now appreciate that MMEJ is a mitotic repair mechanism, but the function of PARP1 in this context remains unclear. Here, we systematically explore the effect that PARP/PARPi have on MMEJ activity in cells with intact NHEJ. Surprisingly, PARP inhibition leads to elevated Polθ-dependent MMEJ levels at ISceI-mediated DSBs, which we find is due to PARP1’s regulation of competing pathways at these breaks. Importantly, we show that PARP is dispensable for MMEJ at double-ended DSBs and is expendable for repair of DSBs during mitosis. Altogether, this data shifts the understanding of PARP’s role in MMEJ and DNA repair pathway choice. Moreover, we believe this data further strengthens a rationale for PARPi/MMEJi combinatorial drug treatment in HR-deficient cancers.

## INTRODUCTION

Double-stranded DNA breaks (DSBs) pose a major threat to genome integrity, as inaccurate DSB repair can result in significant chromosomal aberrations [1]. PARP1 (poly ADP-ribose polymerase 1) is a highly abundant nuclear protein and, among its roles in DNA damage repair [2–4], functions as an initial DSB sensor. DSB-binding activates PARP1, helping orchestrate the DNA damage response (DDR) and repair [3, 5, 6]. PARP1 uses nicotinamide adenine dinucleotide (NAD+) to polymerize poly (ADP)-ribose (PAR) onto many substrates, including itself, chromatin, and other DDR proteins [2, 4, 7]. PARP1, the most extensively studied member within the family of 17 related ADP-ribosyl transferases (ARTDs), accounts for up to 80-90% of PARylation events [8]. However, PARP1, PARP2, and PARP3 have all been described as functioning in response to DNA damage [9].

PARP1-dependent vulnerability was first exploited clinically when it was found that homologous recombination-deficient (HRD) *BRCA1*- and *BRCA2*-null cells are uniquely sensitive to PARP inhibition [10, 11]. Since this seminal work, PARP inhibitors (PARPi) have become a first-line maintenance treatment for HRD cancers [12, 13]. PARPi (e.g., olaparib) functions by competing with NAD+ for binding on both PARP1 and PARP2. PARPi are proposed to be lethal through multiple mechanisms, including PARP trapping, inhibiting PARP catalytic activity, increasing single-stranded DNA (ssDNA) gaps, and destabilizing the replication fork machinery [10, 11, 14–21]. However, only 50% of HRD tumors respond to PARPi treatment due to intrinsic resistance, and, of those that respond, most patients eventually relapse due to acquired drug resistance [22–24]. PARPi resistance in HRD cancer cells can occur through a myriad of mechanisms, including restoration of homologous recombination (HR), DNA replication fork protection, diminished PARP trapping, PARPi efflux, chromatin remodeling, and, in the case of *BRCA*1 mutated cancers, *53BP1* mutation [14, 25–34]. Thus, overcoming PARPi resistance is a key priority and has ushered the development of PARPi combination strategies to enhance treatment efficacy. Current efforts to combat resistance include PARPi combined with drugs that target cell cycle progression, immune checkpoint blockade, angiogenesis, and the DDR pathway [12]. Specifically, one DDR/PARPi combination approach targets the canonical polymerase of microhomology-mediated end joining (MMEJ), Polymerase Q (Polθ, encoded by *POLQ*), as Polθ is essential in HRD cells [35–38]. Notably, Polθ inhibition enhances the cytotoxicity of PARPi and, in some cases, even restores sensitivity to PARPi in PARPi-resistant cells [35, 38]. Currently, clinical trials are assessing the effectiveness of Polθi in combination with PARPi to treat HRD cancers [39]. Thus, understanding the interplay and mechanisms of action between MMEJ and PARP inhibitors is critical.

MMEJ is an intrinsically mutagenic DSB repair pathway. Mammalian MMEJ begins with initial resection at the break-site, in which Mre11/Rad50/Nbs1 (MRN) and CtIP are involved [40, 41]. Resection leads to exposure and annealing of microhomologies flanking the break, allowing for synapsis of the ends through Polθ [39, 42, 43]. Microhomology annealing results in the formation of non-homologous 3’ overhangs that are subsequently removed by nucleases Ape2 [44] or Xpf/Ercc1 [45]. Polθ’s polymerase domain can then use the microhomologies as primers to perform fill-in synthesis [36, 46, 47]. Polδ, Polλ, and Polβ have also been shown to perform this function in varying contexts [43, 48–50]. Finally, the DNA ends are ligated by Lig3-Xrcc1 or Lig1 [51–54]. The essentiality of PARP1 in regulating MMEJ, however, is controversial. PARP1 was first identified to promote mammalian MMEJ through *in vitro* DNA pulldown assays demonstrating that PARP1 can mediate DNA synapsis at overhangs in HeLa cells, independently of the canonical non-homologous end joining (NHEJ) proteins, DNA-PKcs and Ku70/80 heterodimer [55]. Later, the Ku70/80 complex was shown to compete with PARP1 for DNA binding [56]. PARP1’s promotion of MMEJ repair was then further demonstrated using other cell-free methods [57, 58]. In NHEJ-deficient cells, PARP1 promotes MMEJ during class switch recombination [59] and chromosomal translocations [60]. Moreover, at deprotected *XRCC5*^-/-^ (Ku80) mouse telomeres, we and others have observed that MMEJ-dependent telomere fusions are abrogated when treated with PARPi [44, 61]. In mammalian cells, PARPi and PARP knockdown lead to reduced Polθ recruitment to irradiation sites [36, 62]. Altogether, these data suggest that PARP1’s role in MMEJ may be to promote DNA synapsis and to recruit Polθ to DSBs. However, it is important to note that much of the data regarding PARP1’s role in MMEJ was collected in interphase and/or in the absence of canonical NHEJ factors that physiologically compete with MMEJ.

MMEJ strongly competes with NHEJ in G1 and with NHEJ and HR in S/G2 phases [34, 37, 40, 63–66]. Meanwhile, MMEJ is a highly active DSB repair pathway in mitosis, where NHEJ and HR are actively inhibited [67–73]. For example, it has been described that in the absence of *BRCA2*, replication-induced DSBs are repaired by MMEJ in mitosis [74]. Moreover, two recent publications showed that Polθ activity and recruitment to DSBs are highly dependent on mitotic-specific factors [68, 69]. These data suggest that mitosis is possibly the cell cycle phase when MMEJ is most active and physiologically relevant. Considering mitotic-specific recruitment of Polθ to DSBs, it is unclear what role PARP1 plays in MMEJ in cycling cells when all DSB repair pathways are active and competing.

Utilizing a DNA repair reporter, we assessed MMEJ levels following treatment with various PARP inhibitors, including those with minimal PARP trapping activity. Remarkably, PARP inhibition systematically led to elevated *POLQ*-dependent MMEJ levels in cells with intact repair pathways, cells treated with an NHEJ inhibitor, and in HR-deficient (PEO1, *BRCA2*-mutated) cells. We further found that MMEJ upregulation is mediated by inhibition of PARP1, not PARP2, and due to PARP1 promoting NHEJ and HR at certain breaks, likely those with non-blunt ends. Finally, we show that PARP is dispensable for MMEJ at double-ended DSBs and is expendable for repair of DSBs occurring in mitosis. Overall, our findings challenge the current paradigm by demonstrating that PARP does not promote MMEJ. These results have important implications for the development of combination therapies using MMEJ and PARP inhibitors to treat HRD cancers. Additionally, they pave the way for new studies on PARPi resistance mechanisms, which may be directly linked to MMEJ activity.

## MATERIAL AND METHODS

### Cell lines, cell culture, and chemical compounds

Fibrosarcoma HT1080 cells were purchased from ATCC (CCL-121). MEFs TRF2^F/-^ Rosa26-CreERT2 were established in the de Lange lab and obtained from ATCC (CRL-3317). Lenti-X 293T cells were derived from a transformed human embryonic kidney cell line and purchased from Takara (#632180). Cell lines and their derivatives were grown in DMEM medium (Corning #10-013CV) with 10% Cosmic Calf Serum (Cytiva HyClone SH30087.04), 100 mg/mL Penicillin/Streptomycin L-Glutamine (Corning #30-009-CI), and 1X MEM Nonessential Amino Acids (Corning #25-025CI). PEO1 olaparib-resistant (PEO1-OR) cells were created by dose-escalating olaparib in PEO1 (*TP53/BRCA2*-mutated) cells [30] which were obtained from the Gynecologic Tissue and Fluid Bank (GTFB) at the University of Colorado and authenticated at the University of Arizona Genomics Core using short tandem repeat DNA profiling [30]. PEO1-OR cells were maintained in RPMI 1640 supplemented with 10% Cosmic Calf Serum (Cytiva HyClone SH30087.04) and 100 mg/mL Penicillin/Streptomycin L-Glutamine (Corning #30-009-CI). All cell lines were maintained at 3% O_2_ and 7.5% CO_2_.

#### Chemical compounds used include

DMSO (1%, ThermoFisher, #J66650.AP), olaparib (500 nM-20 µM, Selleckchem, AZD2281 #S1060), DNA-PKcsi (1 µM in HT0180 cells and 2.5 µM in PEO1-OR cells, NU7441, R&D Systems #3712/10), and Polθi (10 µM, ART558, MedChemExpress Cat. No.: HY-141520), nocodazole 100 ng/ml (MedChemExpress, #HY-13530), and RO3306 9 µm (MedChemExpress, #HY-12529).Equation

### Plasmids

The MMEJ reporter (pLenti-Puro-MMEJ_rep) used in experiments was described previously [44]. The ISceI-BFP plasmid to cut the MMEJ reporter was derived from the pCBASceI plasmid (Addgene #26477) in which nucleotides 1730-2587 have been replaced by codon optimized ISceI-T2A-BFP. The Cas9-GFP plasmid to cut the MMEJ reporter was derived from pSpCas9(BB)-2A-GFP plasmid (Addgene #48138), and golden gate was used to insert either the non-targeting guide (CTCGACAGTTCGTCCCGAGC) or a guide against the MMEJ reporter (CTGCAAGATTAGGGATAACA). The GFP-PARP1 plasmid used to complement *PARP1*-KO clones was a generous gift from the Karolin Luger lab and was derived by cloning the human *PARP1* coding region into the plasmid pEGFP-C3 (Clontech # 6082-1). The *GFP-PARP1* sequence was then inserted into the lentiviral backbone of pLenti-Puro-MMEJ_rep where the MMEJ reporter was removed from bp 4121 to 5641 and replaced with *GFP-PARP1*, and then Puromycin selection was replaced with Hygromycin B selection from bp 6226 to 6775. Site-directed mutagenesis was then used to mutate the PAM sequence of the plasmid corresponding to the gRNA used to knock out *PARP1*. The *PARP2* plasmid used to complement *PARP2*-KO clones was a generous gift from the Karolin Luger lab and was derived by cloning the human *PARP2* coding region into plasmid mCherry C1 (Addgene, #58476). The *PARP2* sequence was then inserted into the lentiviral backbone of pLenti-Puro-MMEJ_rep where the MMEJ reporter was removed from bp 4121 to 5641 and replaced with *PARP2*, and then puromycin selection was replaced with Hygromycin B selection from bp 6226 to 6775. Site-directed mutagenesis was then used to mutate the PAM sequence on the plasmid corresponding to the gRNA used to knock out *PARP2*. The Cas9 plasmid used to knockout all the genes of interest was derived from TLCV299 (A gift from Adam Karpf, Addgene #87360). This plasmid was modified by removing eGFP and replacing puromycin selection with blasticidin selection. sgRNAs were cloned using golden gate. All plasmids were verified by nanopore sequencing.

### Lentiviral transductions

Lenti-X 293T cells (550,000) were seeded into one well of a 6-well plate. 24 hours later, 2 mL complete DMEM media with 10 mM HEPES was replenished on cells. Cells were then immediately transfected with packaging plasmids (0.5 μg pCMV-VSV-G, 1.6 μg pD8.9), 1.7 μg transfer plasmid, and 11.1 μg of PEI MAX (Polysciences Inc., #24765-1) in 500uL Opti-MEM (Gibco, #31985070). Media was changed after 24 h with 3 mL of complete DMEM. Virus-containing supernatant was collected 48 h and 72 h post-transfection, filter sterilized on a 0.45 mm PES Filter membrane (Whatman Uniflo, #9915-2504) and directly used to infect cells in the presence of 5 mg/ml polybrene (EMD Millipore, #TR-1003-G). Forty-eight hours after infection, cells were washed and selected with 1 mg/ml puromycin (Alfa Aesar, #J61278-MC, 200 mg/ml Hygromycin B (Biosciences, #31282-04-9), or 10 mg/ml blasticidin (RPI, #B12200-0.05). Selection was performed until complete death of uninfected control cells.

### Establishment of knock-out cell lines

For knockout of human *PARP1*, *PARP2*, *EXO1*, *WRN*, and *53BP1*, HT1080 cells were transduced with pLentiGuide-Cas9-Blast and seeded for clones. The single guide RNA sequences for each gene knockout are as follows: PARP1 (CGATGCCTATTACTGCACTG), PARP2 (TTGTTCAGGCAATCTCAACA), EXO1 (TCAGGGGGTAGATTGCCTCG), WRN (ATCCTGTGGAACATACCATG), and *53BP1* (TCATGTGACGATGTAAGACA). The non-targeting guide (CTCGACAGTTCGTCCCGAGC) was included as a Cas9 control. Guide cloning was done following the golden gate cloning protocol and verified by Sanger sequencing. Clonal lines were isolated and clonal knockout verified by immunoblot.

### MMEJ ISceI/Cas9 fluorescent reporter assay

HT1080 cells with or without *PARP1*^-/-^, *PARP2*^-/-^, *EXO1*^-/-^, or *WRN*^-/-^ were transduced with pLenti-Puro_MMEJ_rep and selected with puromycin. Cells were seeded onto 12-well plates 24 h before adding either ISceI or Cas9. Cells were seeded such that they would be 80-90% confluent 24 h later. Then, cells were transfected with either ISceI-T2A-BFP or Cas9-T2A-GFP-guide1. Transfections were done with lipofectamine 3000 (Invitrogen, #L3000015) using 1 µg DNA, 2 µL p3000, and 2 µL lipofectamine 3000 per reaction (following manufacturer’s instructions). 5 h later, cells were split 1/3 into six-well plates and drugs were added. At day 3, cells were analyzed by flow cytometry (MACS Quant VYB, Miltenyi Biotec) for BFP and mCherry expression. BFP+ (ISceI+) or GFP+ (Cas9+) cell populations were gated and mCherry+ cells were quantified.

### Amplicon sequencing of MMEJ reporter

HT1080 cells were transfected with ISceI-T2A-BFP. Five hours later, cells were treated with DMSO (1%), DNA-PKcsi (NU7441, 1 µM), Polθi (ART558, 10 µM), or olaparib (5 µM). At day 3, cells were harvested. Genomic DNA was extracted using QuickExtract (Biosearch Technologies, #QE09050). One PCR reaction was done on each condition using Illumina MiSeq adapter sequences linked to primers that amplify a 78 bp sequence around the ISceI cut region of the reporter. Amplification was done in 50 µL reactions with Q5 High-Fidelity 2X Master Mix (New England Biolabs, #M0492S), 1 µg template, and 0.25 µM of primers for 30 cycles. Products were purified using gel extraction of the bands in 0.6% agarose gel. Using manufacturer instructions, concentration was calculated using TapeStation 4150 (Agilent, #G2992AA). Libraries were pooled at 2.5 nM and sequenced at the University of Colorado Genomics Shared Resource (RRID:SCR_021984). NovaSeq 6000 sequencing system was used, with paired end reads of 2X 150 bp for 50 million reads. Raw sequences were analyzed for DNA repair products at the University of Colorado Anschutz Medical Campus Cancer Center Bioinformatics Core Facility (RRID:SCR_021983). The overlapping forward and reverse raw sequence reads were combined into end-to-end sequences of fragments with NGmerge version 0.3 [cite https://bmcbioinformatics.biomedcentral.com/articles/10.1186/s12859-018-2579-2], allowing a mismatch fraction of 0.1. The adapter trimming software skewer version v0.2.2 [cite https://bmcbioinformatics.biomedcentral.com/articles/10.1186/1471-2105-15-182] was used to remove the sequence TTTCCGGAAGGGTTCAAGTGGGAGAGGGTAATGAATTTTGAGGATGGCGGGGTTGTCACCGTTA from the 5’-end with parameters “-m head -Q 20 -r 0.1 -d 0.05 -n −l 20” and the sequence ACAAAGTGAAGTTGCGAGGAACAAATTTCCCAAGCGA from the 3’-end with parameters “-m tail -Q 20 -r 0.1 -d 0.05 -n −l 10 -L 70” in order to focus the analysis on the relevant part of the fragment sequences. Low-quality sequences were removed using the fastq_quality_filter tool from the FastX toolkit version 0.0.14 [https://github.com/agordon/fastx_toolkit] with parameters “-q 20 -p 100”. Sequences that did not possess the expected 5’- and 3’-ends were filtered out with seqkit grep version 2.6.1 with parameters “-s -r -P -p ‘^GCTT|AGC(?:GA?)?$’”. Identical sequences were enumerated using the fastx_collapser tool from the FastX toolkit.

### Immunoblotting

Cells were collected and washed once using cold PBS. Protein was extracted using RIPA buffer (50 mM Tris-HCl pH 8, 1% NP-40, 0.5% sodium deoxycholate, 0.1% SDS, 150 mM NaCl) combined with protease and phosphatase inhibitor cocktail (Roche, #05892791001) and 1:100 benzonase (Millipore, #70746). Protein concentration was measured using the bicinchoninic acid (BCA) protein assay (Thermo Fisher Scientific, #23227). A total of 25 µg of protein extract was separated on NuPAGE Bis-Tris 4-12% gels (Invitrogen, #WG1402BOX) and transferred onto a PVDF membrane (Cytiva, #GE10600058). Membrane was blocked by shaking 1 h at room temperature in 5% bovine serum albumin (BSA) in PBS-Tween (PBST). The membrane was incubated overnight, rocking at 4°C with primary antibodies diluted in 5% BSA in PBST. The next day, the membrane was washed 3 times with PBST, incubated for 1 h at room temperature in secondary antibody diluted in 5% BSA in PBST, and washed 3 times with PBST. Proteins were detected using ProSignal Fempto (Prometheus, #20-302). Blots were imaged using G:Box chemi XX6 (Syngene).

#### Antibodies used for immunoblotting

PARP1 (diluted 1:2000, Cell Signaling Technology #9542), PARP2 (diluted 1:2000, Active Motif #39744, Antibody has been discontinued), β-Actin (diluted 1:5000, Cell Signaling Technology #8457), WRN (diluted 1:2000, Cell Signaling Technology #4666, RRID:AB_10692114), EXO1 (diluted 1:1000, Cell Signaling Technology #63862), PAR (diluted 1:2000, Cell Signaling Technology #87733), 53BP1 (diluted 1:2000, Novus Biologicals #NB100-304), H3 (diluted 1:5000, Cell Signaling Technology #9715), anti-mouse IgG, HRP-linked (diluted 1:5000, Cell Signaling #7076), anti-rabbit IgG, HRP-linked (diluted 1:5000, Cell Signaling #7074).

### Propidium Iodide (PI) and pH3S10 staining

Protocol from [75] was used. Briefly, cells were trypsinized, spun down, and washed in PBS. Cells were fixed with 70% ethanol and kept at −20°C until use. Cells were spun down, washed in PBS, and resuspended in 1.4 mL PBS with 0.25% Triton X-100 on ice for 15 minutes. Cells were washed and resuspended in 100 μL of PBSBA (PBS containing 1% bovine serum albumin (w/v)) with 0.75 μg anti-phosphorylated histone H3 serine 10 (pH3S10), Alexa Fluor 488 Conjugate (Millipore Sigma, #06-570-AF488) on a rocker, protected from light for 3 h. Cells were then spun down, washed in PBSBA, spun down again, and resuspended in 220 μL propidium iodide (PI) solution (500 μg/mL RNAse (Thermo Fisher Scientific, #EN0531), PI 100 μg/mL in PBS) for 30 minutes at 37°C. Cells were then analyzed by flow cytometry (MACS Quant VYB, Miltenyi Biotec) to identify FITC+ cells and PI+ cells.

### Immunofluorescence of DSBs during mitosis

Cells were seeded on glass coverslips and, once attached, treated with 9 µM RO3306 for 16 hours. After 15-hour treatment, DMSO (1%), olaparib (5 µM), or Polθi (ART558, 10 µM) were added. Cells were then washed with PBS and released into fresh media containing 10 ng/ml nocodazole as well as DMSO (1%), olaparib, or Polθi. Forty minutes after release, cells were irradiated at 2 Gy using a Faxitron Cabinet X-ray System Model RX-650 and placed back into the incubator for 1 or 5 hours. Cells were then fixed with 3.7% formaldehyde in 1x PBS, washed, and permeabilized 10 minutes with KCM buffer (120 mM KCl, 20 mM NaCl, 10 mM Tris pH 7.5, 0.1% Triton), and stored in PBS at 4°C until immunostaining. Cells were blocked 30 minutes at 37°C with ABDIL (20 mM Tris pH 7.5, 2% BSA, 0.2% Fish Gelatin, 150 mM NaCl, 0.1% Triton, 0.1% Sodium Azide), incubated 1.5 hours at 37°C with primary antibodies diluted in ABDIL. Cells were then washed 3 times with PBST and incubated 1 h at room temperature with secondary antibodies, washed again 3 times in PBST and fixed with Fluoroshield containing DAPI (Sigma-Aldrich, #F6057). Imaging was acquired using a NikonTie SDC at 20x magnification through the University of Colorado Light Microscopy Core Facility (RRID:SCR_018993). Image processing and analysis were performed using FIJI (ImageJ) with a custom macro. For each field of view, Z-stacks were acquired, and a sum projection was applied to maximize signal detection. Mitotic cells were identified based on pH3S10 positivity, and regions of interest (ROIs) were manually outlined around mitotic nuclei. Integrated density values for γH2A.X foci were measured within the defined mitotic ROIs.

#### Antibodies used for immunofluorescence (IF)

Phosphorylated γH2A.X (Ser139) (diluted 1:5000, EMD Milipore Cat#05-636), phosphorylated H3S10 (diluted 1:1000, Cell Signaling Technology #9701), goat anti-mouse IgG (H+L) cross-adsorbed secondary antibody, Alexa Fluor 568 (diluted 1:400, ThermoFisher Scientific #A11004), goat anti-rabbit IgG (H+L) cross-adsorbed secondary antibody, Alexa Fluor 488 (diluted 1:400, ThermoFisher Scientific #A11008).

### Analysis of micronuclei

Cells were seeded on glass coverslips and treated with 9 µM RO3306 for 16 hours. After 15 hours of treatment, DMSO (1%), olaparib (5 µM), or Polθi (ART558, 10 µM) were added. Cells were then washed with PBS and released into fresh media containing again DMSO, olaparib, or Polθi. 40 minutes after release, cells were irradiated at 2 Gy using a Faxitron Cabinet X-ray System Model RX-650 and allowed to recover for 5 hours. Cells were then fixed with 3.7% formaldehyde in 1x PBS, washed, and coverslips were mounted with fixed with Fluoroshield containing DAPI (Sigma-Aldrich, Cat#F6057). Cells with or without micronuclei were counted using an Echo Revolve Light Microscope, using a 40x objective. Representative images were taken using the same conditions.

### Statistics

Graphs and statistical analysis were completed in Prism GraphPad (v9). Data are presented as mean ± standard error of the mean (SEM). An unpaired Student’s *t* test was used for statistical comparison between control and treatment groups. A one-way ANOVA was used to determine variance among multiple gestational groups, with a Benjamini-Hochberg multiple comparison post-hoc test to determine significance between individual groups. A *p* value of < 0.05 was considered significant.

## RESULTS

### PARP inhibition increases MMEJ at ISceI-mediated breaks

PARP catalytic activity is described as essential for MMEJ activity, and PARP can be effectively inhibited pharmacologically using PARP inhibitors (PARPi), with olaparib being the most widely studied [76]. Thus, we wanted to examine the MMEJ repair response in the presence of olaparib. To measure MMEJ activity, we used a fluorescent genetic reporter that we previously described [44], consisting of an integrated mCherry cassette with an internal ISceI restriction site flanked by 10 bp microhomologies. The reporter is designed to enable mCherry fluorescence only when cut and repaired by annealing of the 10 bp microhomologies (**Figure 1A**). We transduced the reporter into HT1080 fibrosarcoma cell lines at an MOI of 0.3 to integrate one copy per antibiotic-selected cell. The integrated reporter can then be cut by transiently transfecting a plasmid that expresses ISceI as well as BFP, allowing the identification of the transfected population.

**Figure 1.**
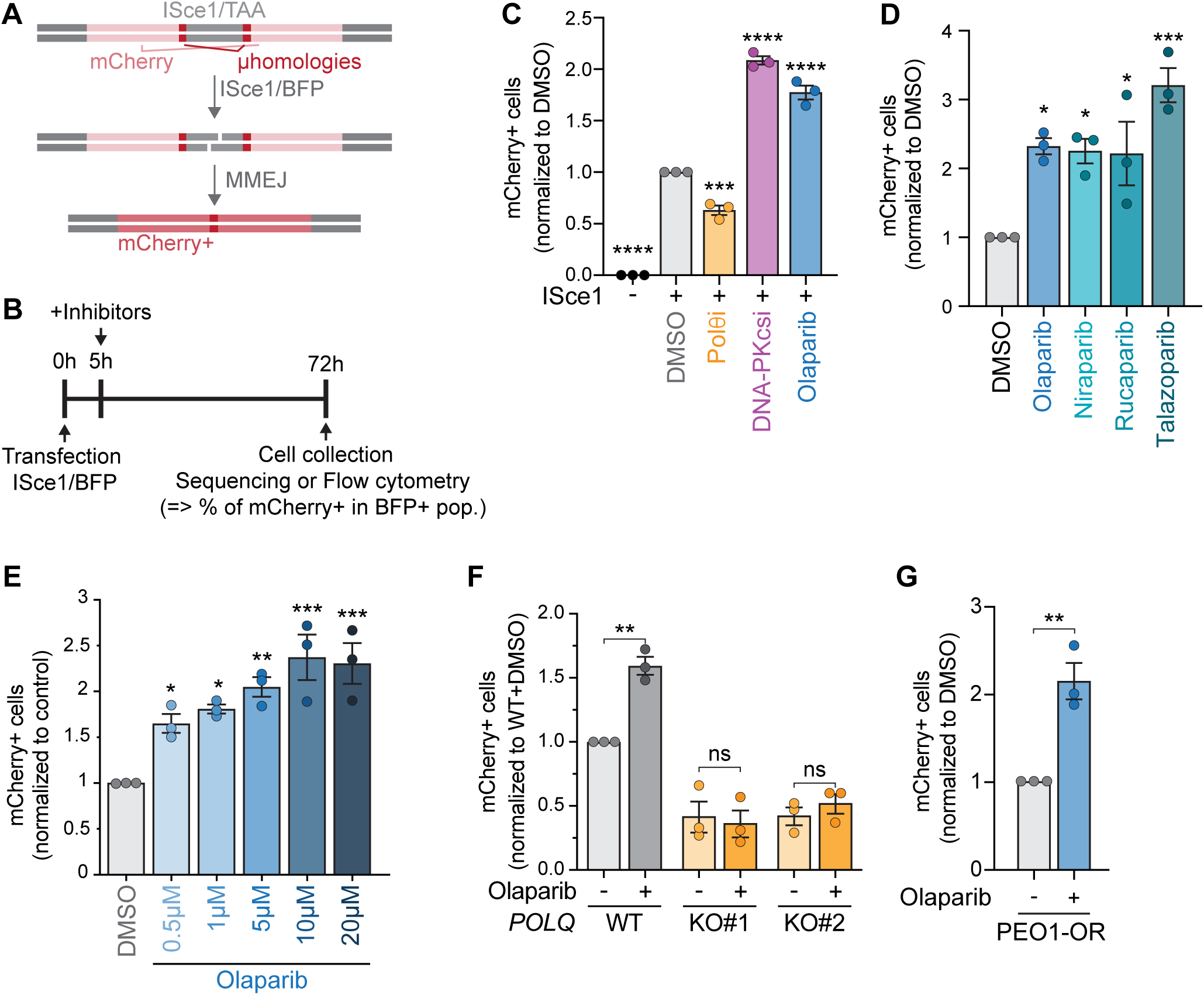
PARP inhibitors increase MMEJ at ISce1-mediated deDSBs. (**A-B**) Schematic (A) and experimental timeline (B) of MMEJ reporter used in (C-G), Figure 2 (C) and (D), Figure 3 (A) and (C) and (F-H), and Figure 5 (C) and (D). Cutting the reporter with ISceI and repair by microhomology annealing to an in-frame, functional mCherry gene. Flow cytometry was used to identify the mCherry+ cells (MMEJ+) within the BFP+ (ISceI+) population. (**C-E**) MMEJ quantification using reporter and timeline from (A-B) in HT1080 cells. Values are normalized to DMSO. Drugs used were olaparib (5 µm or as indicated), DNA-PKcsi (NU7441, 1 µm), Polθi (ART558 10 µm), niraparib (2.5 µm), rucaparib (2.5 µm), and talazaparib (2.5 µm). (F) MMEJ quantification in HT1080 parental cells compared to isogenic POLQ-KO cells following olaparib treatment. Values are normalized to wild-type DMSO. (**G**) MMEJ quantification in PEO1-OR cells following olaparib treatment. Values are normalized to DMSO. Statistical analyses for (**C-G**): Data represent three independent experiments, each the average of three technical replicates. Data are mean ± SEM. Statistical test, one way ANOVA with multiple comparison correction. ns: non-significant, *p<0.05, **p<0.01, ***p<0.001, ****p<0.0001.

Five hours after ISceI transfection, we treated cells with PARP, Polθ, or DNA-PKcs inhibitors, and analyzed them by flow cytometry three days later (Experimental timeline depicted in **Figure 1B**). The frequency of mCherry+ cells within the BFP+ population was used to measure MMEJ activity. Importantly, all conditions had similar levels of BFP+ cells, indicating that transfection levels were similar in all conditions (**Supplementary Figure 1A**). As expected, we found no mCherry-positive cells in the control population not transfected with ISceI (**Figure 1C**). Likewise, inhibiting the primary MMEJ polymerase, Polθ (Polθ inhibitor ART558) [35] led to decreased MMEJ activity, while inhibition of the competing pathway, NHEJ (DNA-PKcs inhibitor NU7441), increased MMEJ activity (**Figure 1C**). Finally, we treated cells with 5 μM olaparib, the lowest dose at which PARylation was fully ablated (**Supplementary Figure 1B**) and which is a physiological dose [77]. Strikingly, we found a significant increase in MMEJ, a surprising result considering that PARP1 is described to promote MMEJ (**Figure 1C**). To confirm this result, we complemented the flow-based assays with amplicon sequencing and found higher percentages of MMEJ repair products in olaparib-treated cells compared to untreated cells (**Supplementary Figure 1C**). We also verified that the increase in MMEJ was not specific to olaparib and could be observed with other PARP inhibitors. Indeed, the results showed elevated MMEJ upon treatment with all the PARP inhibitors we tested (**Figure 1D and Supplementary Figure 1D**).

Since MMEJ levels vary over the cell cycle [40, 78], we tested whether PARPi led to cell cycle perturbation. As previously reported [79], olaparib treatment induced a dose-dependent increase in S and G2 phases and a decrease in mitotic index (**Supplementary Figure 1 E-F**). We, therefore, tested escalating doses of olaparib in our MMEJ reporter and found that all tested doses increased MMEJ levels, independently of their effect on the cell cycle (**Figure 1E**). These data suggest that the observed increase in MMEJ is not a consequence of cell cycle perturbation. Of note, it is possible that the effect of higher doses of olaparib on MMEJ would be more prominent if the experimental timeline accounted for population doubling differences.

### PARPi-mediated MMEJ increase is dependent on Polθ and is also observed in HR-deficient cells

While Polθ is the primary polymerase responsible for MMEJ repair [36, 37, 80, 81], data show that other polymerases, such as Polλ [49], can also perform MMEJ repair. Thus, we wanted to determine whether the observed increase in MMEJ upon PARPi treatment depends on the canonical MMEJ polymerase. We measured MMEJ repair in olaparib-treated wild-type and isogenic *POLQ*-KO HT1080 cells [44]. As expected, *POLQ*-KO cells had significantly lower MMEJ levels than wild-type cells (**Figure 1F**). Furthermore, we found that olaparib-mediated effects on MMEJ were abrogated upon *POLQ*-KO, demonstrating that the PARPi-dependent increase in MMEJ is *PolQ*-dependent. Similarly, a pharmacologic approach to inhibit Polθ activity attenuated the PARPi-dependent increase of MMEJ levels (**Supplementary Figure 1G**), indicating that MMEJ increases seen upon olaparib treatment depend on the catalytic activity of Polθ.

An increase in MMEJ upon treatment with PARP inhibitors has broad consequences for the use of PARP inhibitors in patients. Notably, *BRCA2* reversal mutations in patients treated with PARP inhibitors are likely MMEJ-dependent [28, 29]. Thus, one can speculate that PARP inhibition could cause an upregulation of MMEJ that promotes the acquisition of drug resistance. To determine whether the increase in MMEJ also happens in HR-deficient cells, we transduced our reporter into PEO1 ovarian cancer cells, in which *BRCA2* is mutated. Since these cells are highly sensitive to PARP inhibition, we used cells that were made resistant to olaparib through months of drug escalation (PEO1-OR) [30]. We found that *BRCA2* mutated PEO1-OR cells also display increased MMEJ levels upon treatment with olaparib (**Figure 1G**).

In summary, our data show that inhibition of PARP causes an increase in ISceI-induced MMEJ that is dependent on Polθ and independent of *BRCA2*.

### PARPi-mediated increases in MMEJ are dependent on PARP1, not PARP2

Olaparib targets both PARP1 and PARP2 [82, 83]. To gain more mechanistic insight into what causes the MMEJ increase upon olaparib treatment, we wanted to determine whether it was due to inhibition of PARP1, PARP2, or both. We established isogenic *PARP1*-KO and *PARP2*-KO HT1080 clonal cell lines and measured the effect of olaparib on MMEJ. *PARP1* or *PARP2* knockouts were validated via immunoblot (**Figure 2A-B**). As expected, *PARP1*-KO cells, similarly to olaparib-treated cells, showed almost complete ablation of PARylation (**Figure 2A**). Next, we quantified MMEJ levels of *PARP1*-KO and *PARP2*-KO cells upon olaparib treatment. Compared to wild-type cells, olaparib treatment failed to increase MMEJ levels in *PARP1*-KO clones, demonstrating that PARP1 is required to mediate the olaparib-dependent increases in MMEJ levels (**Figure 2C**). Conversely, *PARP2*-KO cells showed increased MMEJ levels with olaparib treatment, similarly to wild-type cells (**Figure 2D**). Altogether, this data implicates catalytic inhibition of PARP1, not PARP2, in specifically modulating DNA repair towards MMEJ.

**Figure 2.**
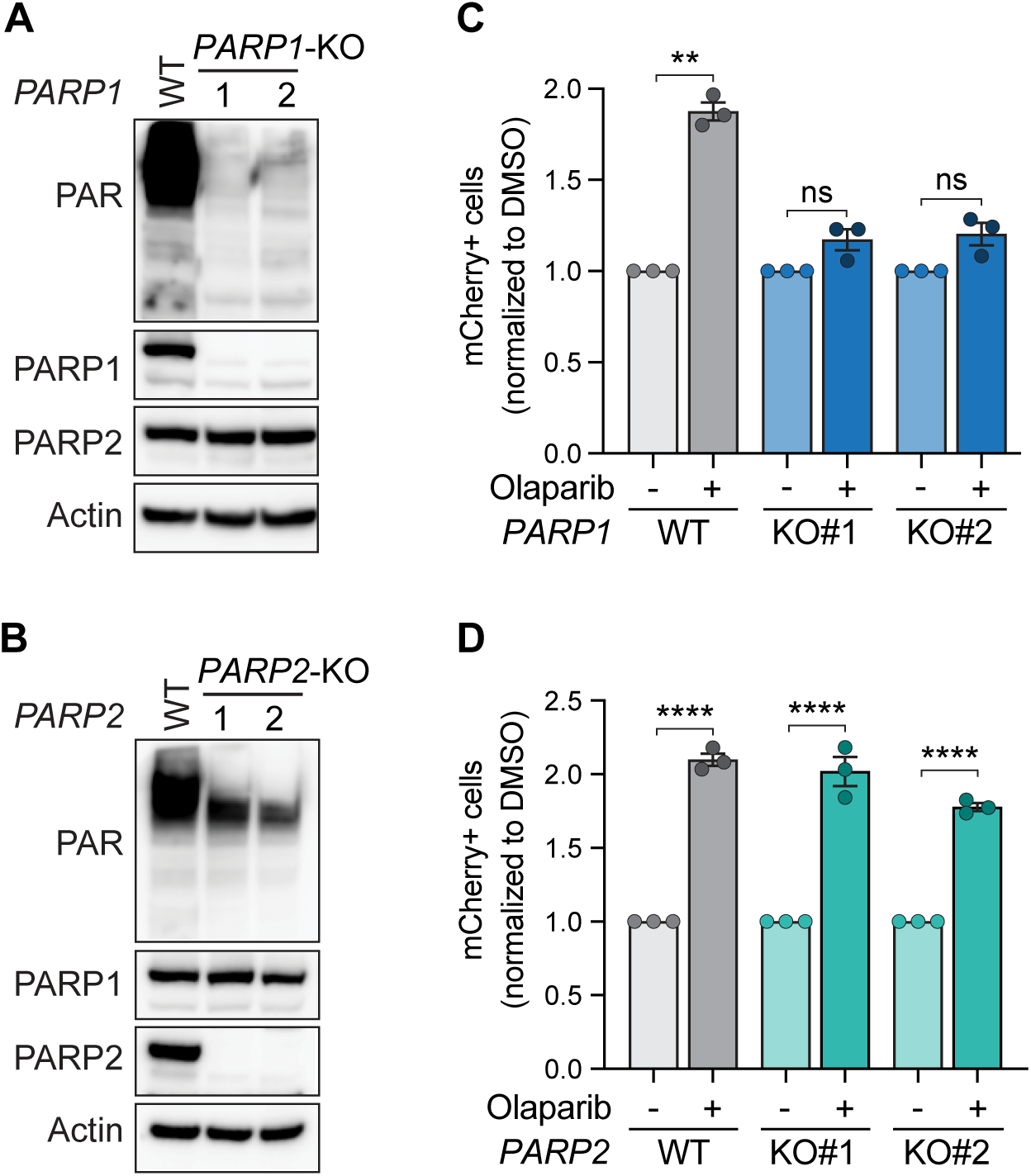
PARPi-mediated MMEJ increase is dependent on PARP1, not PARP2. (**A-B**) Immunoblot of Actin, PARP1, PARP2, and PAR in parental HT1080 cells and isogenic PARP1-KO (A) and PARP2-KO (B) clones. (**C-D**) MMEJ quantification using reporter and experimental timeline from Figure 1A-B in HT1080 parental cells and indicated isogenic PARP1-KO (C) or PARP2-KO clones (D). Values are normalized to each cell line DMSO. Drug used was olaparib (5 µm). Statistical analyses for (**C, D**): Data represent three independent experiments, each the average of three technical replicates. Data are mean ± SEM. Statistical test, one way ANOVA with multiple comparison correction. ns: non-significant, **p<0.01, ****p<0.0001.

### PARPi-mediated MMEJ increase depends on both NHEJ and HR

Next, we sought to determine the mechanism by which PARPi can lead to increased MMEJ following ISceI transfection. It is unlikely that PARP1 directly inhibits MMEJ activity. Instead, we hypothesized that the observed increase in MMEJ is due to the suppression of a competing repair mechanism upon PARP1 suppression. Since olaparib treatment increased MMEJ in *BRCA2*-mutated cells (**Figure 1G**), the increase in MMEJ is unlikely to be due to the suppression of HR alone. On the other hand, PARPi and DNA-PKcsi have a similar effect on MMEJ (**Figure 1C**), suggesting that PARP1 could be promoting NHEJ. We, therefore, tested whether DNA-PKcs inhibition and olaparib were epistatic but found, instead, that the drug combination has an additive effect on MMEJ (**Figure 3A**).

**Figure 3.**
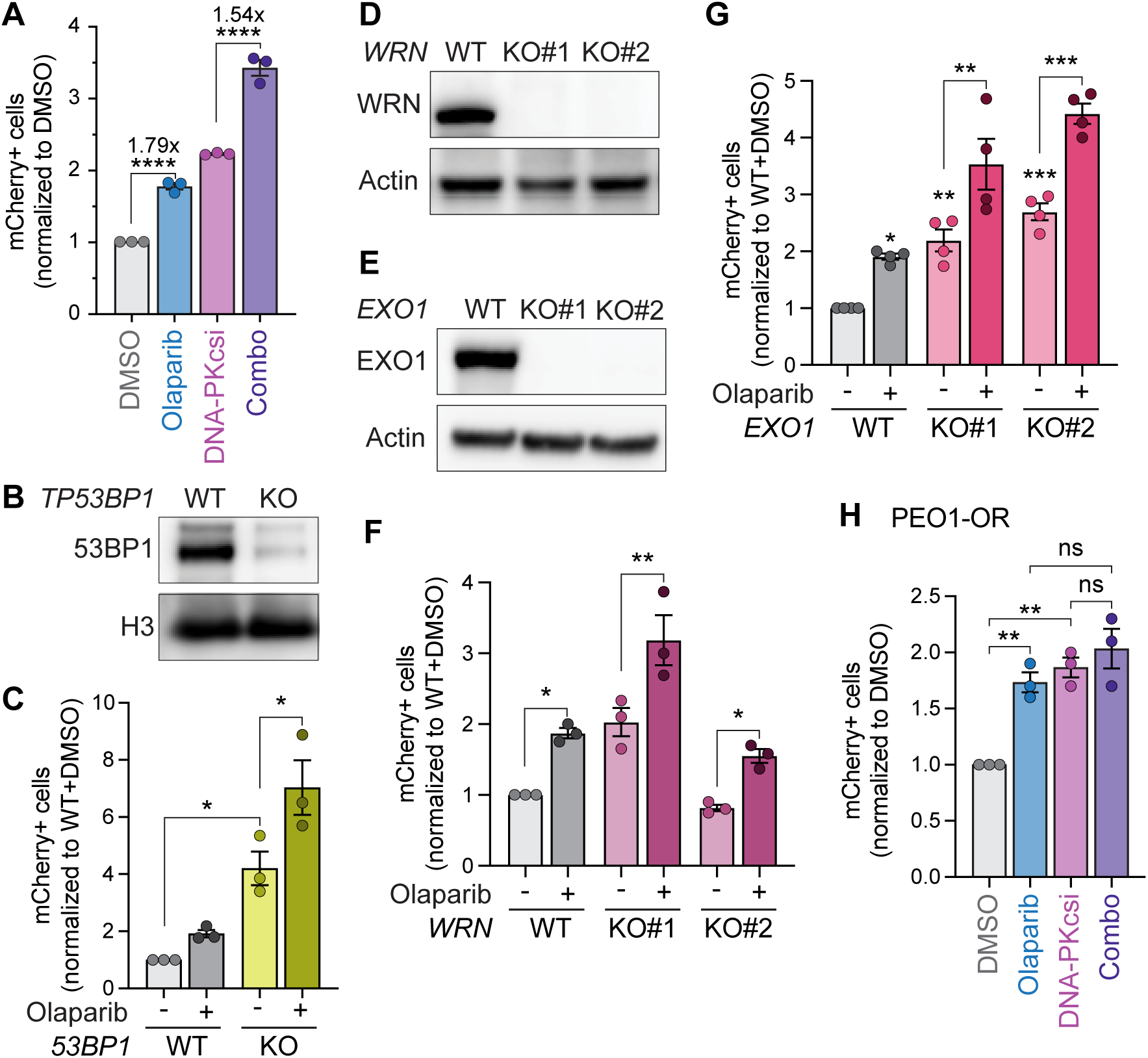
PARPi-dependent MMEJ increase depends on both NHEJ and MMEJ. (**A**) MMEJ quantification using reporter and experimental timeline from Figure 1A-B in HT1080 cells following olaparib (5 µm), DNA-PKcsi (NU7441, 1 µm), or both. Values are normalized to each cell line DMSO. (**B**) Immunoblot of 53BP1 and histone H3 in parental HT1080 cells and isogenic *TP53BP1*-KO cells. (**C**) MMEJ quantification in HT1080 parental cells and indicated isogenic *TP53BP1*-KO cells. Values are normalized to wild-type DMSO. Drug used was olaparib (5 µm). (**D-E**) Immunoblot of Actin, WRN (D), and EXO1 (E) in parental HT1080 cells and isogenic *WRN*-KO (D) and *EXO1*-KO (E) cells. (**F-G**) MMEJ quantification in HT1080 parental cells and indicated isogenic *WRN*-KO (F) or *EXO1*-KO clones (G). Values are normalized to wild-type DMSO. Drug used was olaparib (5 µm). (H) MMEJ quantification in PEO1-OR cells following olaparib (5 µm), DNA-PKcsi (NU7441, 2.5 µm), or combo. Statistical analyses for (**A, C, F-H**). Data represent three (A, C, F, H) or four (G) independent experiments, each the average of three technical replicates. Data are mean ± SEM. Statistical test, one way ANOVA with multiple comparison correction. ns: non-significant, *p<0.05, **p<0.01, ***p<0.001, ****p<0.0001.

When analyzing the sequencing of the ISceI reporter, we noticed that another repair outcome was strongly suppressed by Polθ inhibition and with an MMEJ repair signature (**Supplementary Figure 2**). Noticeably, olaparib had no effect on this repair outcome, which utilizes microhomologies closer to the break site, likely requiring less resection. Thus, we reasoned that PARP1 might regulate break resection, an essential branchpoint in determining DSB repair pathway choice [1, 71]. Inhibition of end resection promotes NHEJ, while short-range resection drives repair towards HR and MMEJ, with HR requiring further long-range resection [39, 71]. The function of PARP in resection remains debated, as it has been shown to both promote and antagonize it [84–86]. First, to test the role of PARP1 at the first resection branchpoint, we knocked out *53BP1*, which inhibits short-range resection [87]. As expected, *53BP1*-KO cells had elevated MMEJ levels. However, olaparib treatment still increased MMEJ in this context (**Figure 3B-C**). Next, we set to determine whether PARP1 could promote long-range resection that inhibits MMEJ. EXO1 and DNA2 are nucleases responsible for long-range resection leading to HR [88–90]. While DNA2 is essential in the experimental cell line, it can form complexes with WRN helicase to perform its function [91, 92]. Hence, we created isogenic *WRN*-KO and *EXO1*-KO cells and measured MMEJ levels in response to olaparib (**Figure 3D-G**). Notably, *EXO1*-KO showed a baseline MMEJ higher than wild-type cells, consistent with its role in competing with MMEJ for repair [88] (**Figure 3G**). However, both *WRN*- and *EXO1*-KO cell lines still showed increased MMEJ levels following olaparib treatment (**Figure 3F-G**).

Since no single factor could be identified as the primary driver of MMEJ increase, we reasoned that this effect likely results from multiple functions of PARP1 in DSB repair. To test this, we measured MMEJ following the dual inhibition of both NHEJ and HR. Specifically, we inhibited DNA-PKcs in *BRCA2*-mutated PEO1-OR cells and quantified MMEJ after additional olaparib treatment. While inhibition of PARP and DNA-PKcs significantly increased MMEJ levels individually, the combined treatment had no additional effect (**Figure 3H**). Thus, when both HR and NHEJ are suppressed, PARP inhibition does not further increase MMEJ. Together, these findings suggest that PARP1 inhibition suppresses multiple repair mechanisms, leaving some DSBs unrepaired by NHEJ and HR, ultimately diverting them toward MMEJ.

### PARPi-dependent MMEJ increase does not occur at Cas9-induced blunt breaks

Finally, we wanted to determine whether PARPi increases MMEJ at different DSB repair substrates. Cleavage by ISceI leaves 4 nucleotide 3’ overhangs (**Supplementary Figure 2B**). To generate blunt ends, we induced DSBs using Cas9 instead of ISceI (**Figure 4A**). Using the same experimental timeline as with the ISceI system, we transiently transfected a plasmid expressing Cas9, a guide RNA targeting our reporter, and GFP (**Figure 4B**). As expected, no fluorescence was detected without Cas9 transfection (**Figure 4C**). Meanwhile, upon Cas9 DSB induction, the percentage of mCherry+ cells decreased upon Polθ inhibition and increased upon DNA-PKcs inhibition (**Figure 4C**). Interestingly, olaparib led to no measurable difference in Cas9-mediated MMEJ levels (**Figure 4C**), a result that remained consistent across a range of olaparib doses (**Figure 4D**). Similarly, double inhibition of PARP and Polθ had no effect compared to Polθi only (**Figure 4E**). While PARP inhibition has different effects on specific MMEJ repair outcomes of the same break type (**Supplementary Figure 2A**), these data demonstrate that PARP also has differential effects on the repair of blunt end breaks compared to those with small 3’ overhangs.

**Figure 4.**
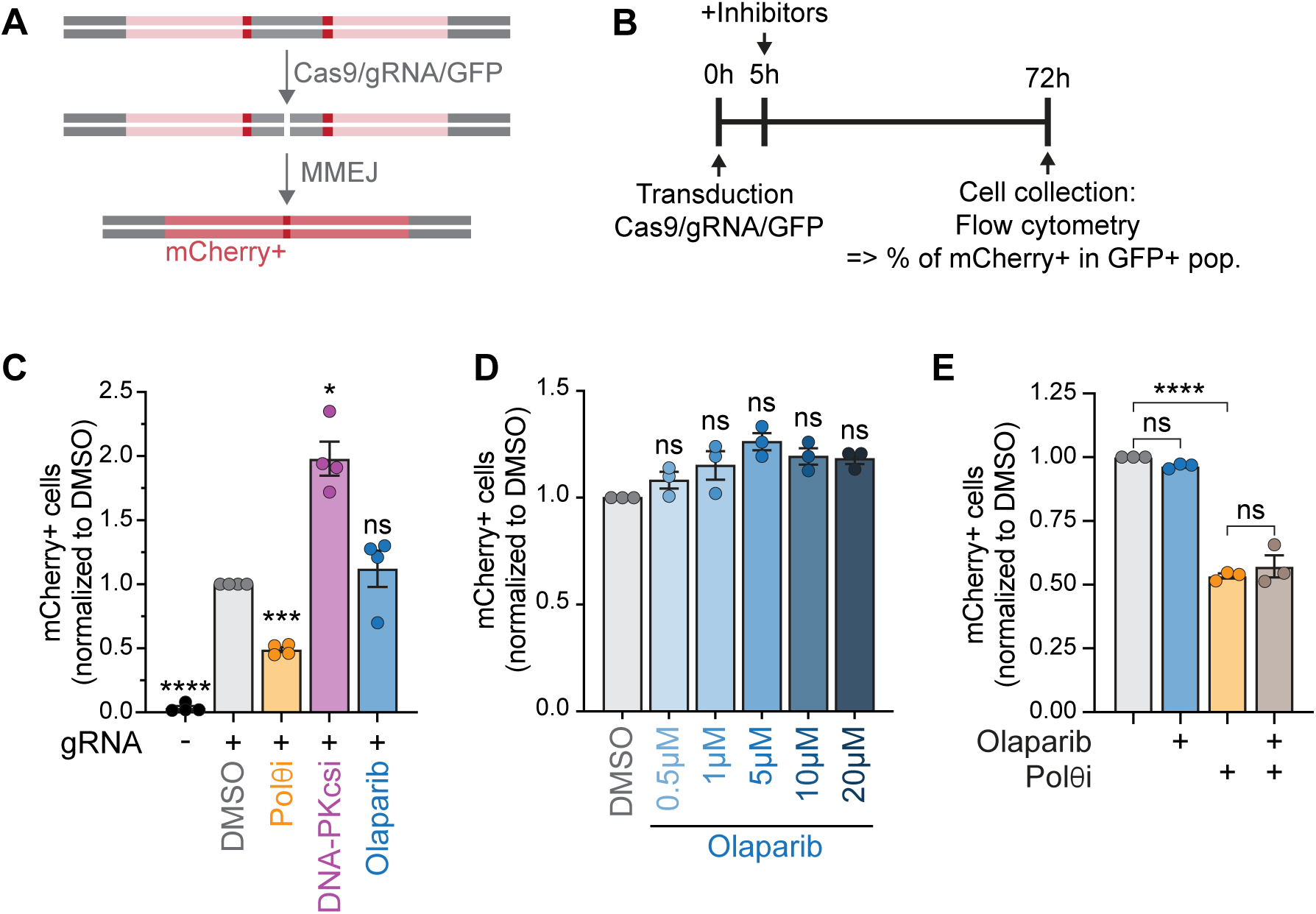
PARPi-dependent MMEJ increase does not occur at Cas9-induced blunt DSBs. (**A-B**) Schematic (A) and experimental timeline (B) of MMEJ reporter used in (C-E) and Figure 5 (E). Repair of the Cas9-induced cut with microhomology annealing leads to an in-frame, functional mCherry gene. Flow cytometry was used to identify the mCherry+ cells (MMEJ+) within the GFP+ (Cas9+) population. (**C-E**) MMEJ quantification using the reporter and timeline from (A-B) in HT1080 cells. Values are normalized to DMSO. Drugs used were olaparib (5 µm or indicated dose), DNA-PKcsi (NU7441, 1 µm), Polθi (ART558, 10 µm). Statistical analyses (**C-E**): Data represent three (D, E) or four (C) independent experiments, each the average of three technical replicates. Data are mean ± SEM. Statistical test, one way ANOVA with multiple comparison correction. ns: non-significant, *p<0.05, ***p<0.001, ****p<0.0001.

Overall, these data suggest that PARP1 could regulate factors involved in the processing of dirty DSBs, ultimately determining whether MMEJ repair occurs. Nonetheless, in all the experimental settings tested, PARP inhibition never decreased MMEJ levels, suggesting that PARP is dispensable for MMEJ repair of intrachromosomal breaks.

### PARP1 and PARP2 are dispensable for MMEJ repair at deDSBs

PARP1 was originally identified as a key factor in MMEJ and is widely considered a central promoter of this pathway [39, 71, 78]. Consistently, we and others have shown that PARP inhibition and PARP1 knockdown reduce MMEJ following telomere deprotection in mouse embryonic fibroblast (MEF) cells [44, 61]. However, our data challenge this well-established view, suggesting that PARP may be dispensable for MMEJ, at least in the context of double-ended (de)DSBs in human cells. Given this contradiction with the prevailing model of MMEJ, we sought to rigorously test whether PARP1 and/or PARP2 are truly dispensable for MMEJ repair of blunt or overhang breaks. To this end, we complemented *PARP1*-KO and *PARP2*-KO clones with exogenous GFP-PARP1 and PARP2, respectively (**Figure 5A-B**). While the *PARP2*-KO clones easily re-expressed PARP2 (**Figure 5B**), GFP-PARP1 expression in *PARP1*-KO clones was variable and below wild-type levels (**Figure 5A**). Similarly, PARylation was not fully restored, mirroring GPF-PARP1 protein expression (**Figure 5A**). To restrict our analysis to cells with higher PARP1 re-expression, we took advantage of the fact that PARP1 was fused to GFP and gated our flow cytometry analysis of MMEJ on cells with “high” GFP expression. We compared MMEJ levels between *PARP1* or PARP*2* knock-out and complemented cells following ISceI transfection. Consistent with the data showing that PARPi increases MMEJ (**Figure 1C**), *PARP1*-KO clones showed elevated levels of MMEJ compared to control cells, which were rescued by exogenous PARP1 expression (**Figure 5C**). Notably, the cell cycle of these clones remains unaltered (**Supplementary Figure 3A-B**), further confirming that the observed increase in MMEJ upon PARP suppression is not a consequence of cell cycle perturbations. In contrast, *PARP2*-KO clones showed similar MMEJ levels to control cells, and complementation with exogenous PARP2 did not have any effect (**Figure 5D**). These data parallel our finding that the absence of PARP2 does not change olaparib’s effect on MMEJ (**Figure 2D**).

**Figure 5.**
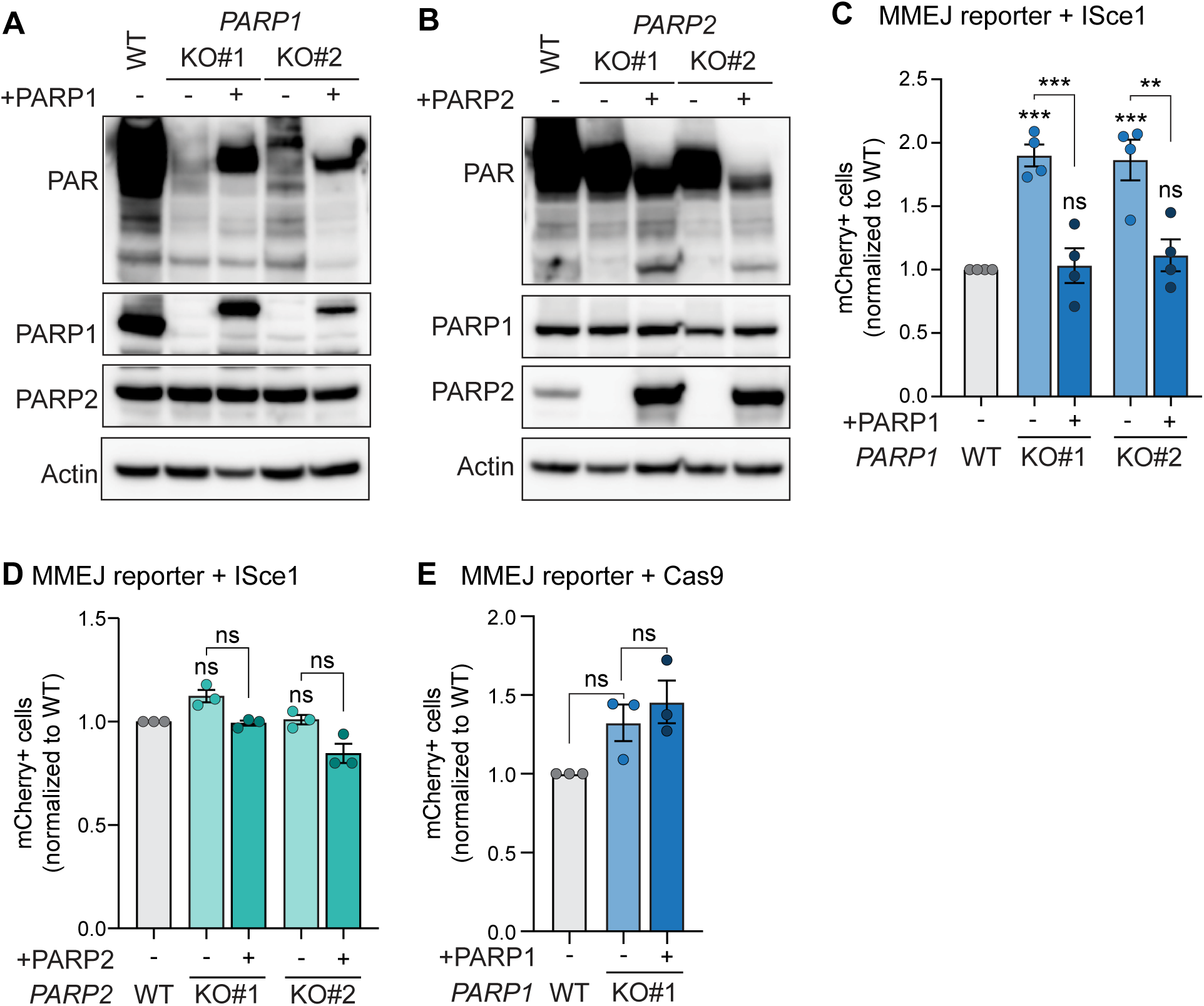
PARP1 and PARP2 are dispensable for MMEJ at deDSBs. (**A-B**) Immunoblot of Actin, PARP1, PARP2, and PAR in HT1080 cells and isogenic *PARP1*-KO with or without complemented GFP-PARP1 (A) or *PARP2*-KO with or without complemented PARP2 (B). (**C-D**) MMEJ quantification using reporter with ISceI cut (from Figure 1A-B) in HT1080 parental cells and indicated isogenic *PARP1*-KO with or without complemented GFP-PARP1 (C) or *PARP2*-KO with or without complemented PARP2 (D). Values are normalized to wild-type DMSO. Drug used was olaparib (5 µm). (**E**) MMEJ quantification using reporter with Cas9 cut (from Figure 4A-B) in HT1080 cells and isogenic *PARP1*-KO cells with or without complemented GFP-PARP1. Values are normalized to wild-type DMSO. Statistical analyses (**C-E**): Data represent four (C) or three independent (D-E) experiments, each the average of three technical replicates. Data are mean ± SEM. Statistical test, one way ANOVA with multiple comparison correction. ns: non-significant, **p<0.01, ***p<0.001.

Finally, we assessed the effect of *PARP1* and *PARP2* knockout complementation on Cas9-induced MMEJ. Since both the PARP1 and Cas9 plasmids expressed GFP, we could not analyze mCherry+ cells in the GFP+ population in this experiment; instead, we measured the total mCherry+ population. Similarly, we could not overcome the low expression of PARP1 in clone 2 by analyzing specifically the high Cas9-GFP+ cells. Therefore, we limited our analysis to clone 1. Consistent with our previous data, we found that neither *PARP1* knockout nor its rescue influenced MMEJ at Cas9-induced DSBs (**Figure 5E**). Similarly, MMEJ levels remained unchanged in *PARP2*-KO and *PARP2*-KO complemented cells (**Supplementary Figure 3C**). Altogether, our data demonstrate that PARP1 and PARP2 are dispensable for repair of both blunt and overhang deDSBs in human cells.

### PARP is dispensable for mitotic DSB repair

*In vitro* data has implicated a role for PARP1 in promoting MMEJ during interphase by competing with KU70/80 for DNA binding, facilitating synapsis formation at the break-site, and recruiting Polθ to the break-site [36, 55–58, 62]. While MMEJ is active in all cell cycle phases, it competes with NHEJ during G1 and with NHEJ and HR in S and G2 [34, 37, 40, 63–66]. Therefore, in cells proficient for NHEJ and HR, low levels of MMEJ activity are expected during interphase. Conversely, two recent studies have shown that MMEJ is active during mitosis, when both NHEJ and HR are suppressed [68, 69]. At this stage, mitotic-specific factors RHINO and PLK1 promote the recruitment of Polθ to DSBs [68, 69]. The observed MMEJ activity measured by our reporter assay is thus likely to occur during mitosis at DSBs that failed to repair through canonical pathways during interphase. Thus, PARP1 may be dispensable for MMEJ specifically in mitosis, when Polθ relies instead on PLK1/RHINO for its recruitment to DSBs. We therefore tested whether PARP is required for the repair of DSBs in mitosis.

Maintaining cells in mitosis for the duration of our reporter experiment is not feasible, as prolonged mitotic arrest leads to telomere deprotection and cell death [93]. Instead, we assessed DSB repair in mitosis by staining for gH2A.X in phosphorylated Histone H3 serine 10+ (pH3S10+) cells following irradiation, as performed in [68]. Cells were arrested at the G2/M border using a CDK1 inhibitor (RO3306) for 16 hours, then released and maintained in mitosis with nocodazole treatment (**Figure 6A**). One hour prior to mitosis release, cells were treated with Polθi or olaparib, and the drugs were maintained thereafter. Forty minutes after RO3306 release, when most cells had reached mitosis, cells were irradiated (2 Gy), then fixed for immunofluorescence 1 and 5 hours later (**Figure 6A**). As previously described [68], gH2A.X foci significantly decreased 5 hours post-irradiation in control cells, signifying resolution of IR-induced damage (**Figure B-C**). However, gH2A.X foci persist in Polθi-treated cells, indicating that mitotic DSB repair relies on Polθ (**Figure 6B-C**). In contrast, a significant decrease in gH2A.X foci were observed in the olaparib-treated cells, indicating that mitotic repair can occur in the absence of PARP activity. To confirm these results, we performed a similar experiment in which cells were not treated with nocodazole but instead allowed to reach G1. We then quantified the number of micronuclei, which arise from unrepaired DSBs in mitosis [94] (**Figure 6D**). As previously reported, Polθ inhibition increased the number of micronuclei in G1 following irradiation [68]. Conversely, treatment with olaparib had no effect on the number of observed micronuclei (**Figure 6D-E**).

**Figure 6.**
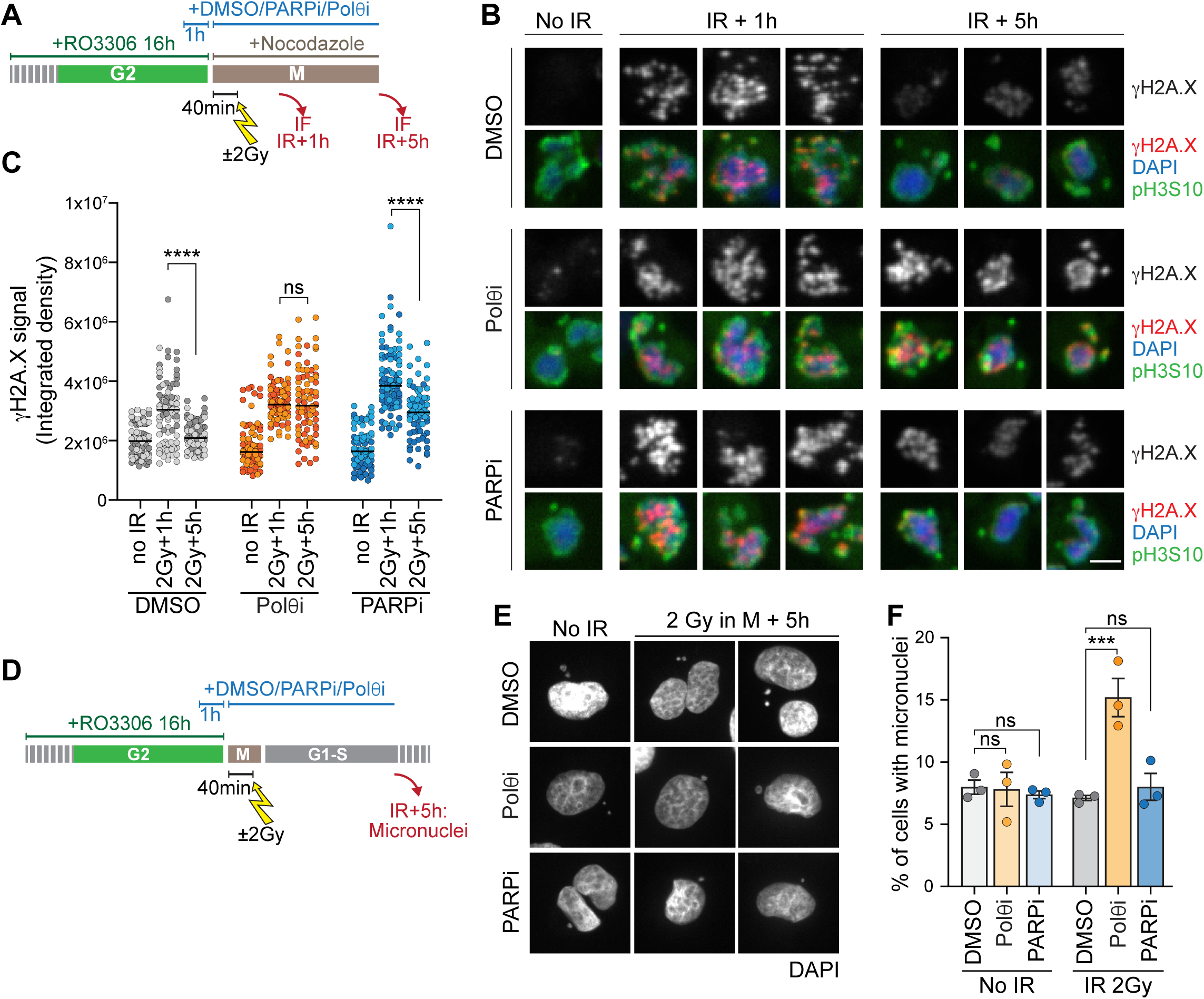
PARP is dispensable for mitotic DSB repair. (**A**) Experimental timeline of (B) and (C). HT1080 cells were arrested in RO3306 (9 µm) for 16 hours. DMSO (1%), olaparib (5 µM), or Polθi (ART558, 10 µM) were added 15 hours after RO3306. After 16 hours in RO3306, cells were released into nocodazole (10 ng/ml) and maintained in respective DMSO, olaparib, or Polθi treatment. Forty minutes after RO3306 wash, cells were irradiated (2 Gy), and samples collected 1 hour and 5 hours later. (**B-C**) Representative images (B) and quantification (C) of IF from HT1080 cells from (A) at 1 hour and 5 hours post irradiation compared to no irradiation. Cells were stained for pH3S10, γH2A.X, and DAPI. Scale bar: 10µm. γH2A.X intensity was quantified in pH3S10+ cells. (**D**) Experimental timeline for (E) and (F). HT1080 cells are arrested in RO3306 (9 µm) for 16 hours. DMSO (1%), olaparib (5 µM), or Polθi (ART558, 10 µM) were added 15 hours after RO3306. After 16 hours in RO3306, RO3306 was removed and cells were maintained in respective DMSO, olaparib, or Polθi treatment. Cells were irradiated (2 Gy) forty minutes after RO3306 release and fixed for micronuclei staining 5 hours later. (**E-F**) Representative images (E) and quantification (F) of IF from HT1080 cells from (D) in cells 5 hours post irradiation compared to no irradiation. Cells were stained for DAPI and the percentage of cells with micronuclei was quantified. Statistical analyses for (**C, F**): Data represent two independent (C) and three independent (F) experiments. For each replicate, at least 35 cells (C) and 300 cells (F) were analyzed. Statistical test, one way ANOVA with multiple comparison correction. ns: non-significant, ***p<0.001, ****p<0.0001.

Altogether, these data demonstrate that PARP is indeed dispensable for DSB repair during mitosis, when MMEJ is the sole active pathway. Hence, we concluded that PARP is dispensable for mitotic MMEJ.

## DISCUSSION

Through mechanistic and functional assays, we demonstrate that PARP is dispensable for MMEJ repair of deDSBs, especially mitotic MMEJ. Initial studies suggested that MMEJ activity depends on PARP1 [55–61, 95]. However, recent data indicate that certain alternative end-joining mechanisms involving PARP1 function independently of Polθ [95, 96]. Consistent with our findings, recent studies also suggest that PARP1 may not be essential for Polθ-dependent MMEJ [97, 98]. When genome-wide breaks were induced at sites with different chromatin contexts, PARP1 primarily promoted MMEJ repair at heterochromatic sites, suggesting that chromatin context influences its role in MMEJ [97]. Additionally, G1-specific DSBs in NHEJ-deficient cells have been observed to undergo repair by Polθ in the S-G2/M phase without requiring PARP1 [98].

Though we cannot be certain the cell cycle phase when MMEJ primarily occurs with our reporter, we used cells that have intact NHEJ and HR pathways, which likely outcompete MMEJ in interphase. Indeed, Ku70 was shown to suppress MMEJ during G1 [96], and we found that MMEJ activity at our reporter is almost null in serum-starved cells (data not shown). On the other hand, Polθ competes with both RPA and RAD51 during S/G2 [36, 99]. Therefore, we speculate that most MMEJ repair at our reporter is likely to occur during mitosis, when HR and NHEJ are suppressed. During this phase of the cell cycle, PLK1-activated RHINO recruits Polθ to DSBs [68, 69], a function previously attributed to PARP1 [36, 62].

Conversely, the earlier studies that described PARP1 as essential to promoting MMEJ were done in the context of DNA-PK suppression, in which MMEJ is likely reactivated in G0/G1 [56, 96, 100]. These include fusions of deprotected telomeres in Ku80-KO MEFs, which occur in G1 [36, 44]. Therefore, we propose the following model: In cells with intact pathways, MMEJ repairs deDSBs mostly during mitosis, when activated RHINO can recruit Polθ, rendering PARP1 dispensable. In cells absent of Ku70, on the other hand, MMEJ is reactivated during G0/G1 [96], where it depends on PARP1 for the recruitment of Polθ. This model suggests that the factors required for MMEJ may vary depending on the cell-cycle context. Since PARP1 was shown to recruit XRCC1 to breaks [5, 55, 101], it would be interesting to determine whether mitotic MMEJ uses XRCC1-LIG3 for the ligation step or, on the other hand, relies on LIG1. Likewise, Polθ is a target for PARylation [99, 102]. It would be interesting to tease apart the function of this PARylation during different phases of the cell cycle.

Similarly, most DSBs in unchallenged cells arise from collapsed replication forks. Thus, single-ended DSBs are likely more physiological than deDSBs, especially in the context of PARP inhibitors. It will be critical to determine which factors promote MMEJ at these types of breaks.

We also found that PARPi treatment led to an increase in MMEJ levels following ISceI-mediated intrachromosomal breaks. This increase was specifically dependent on PARP1, not PARP2. These findings illustrate that PARP1 and PARP2 have distinct effects on MMEJ, likely due to differential roles in competing pathways. Indeed, several labs have reported divergent functions between PARP1 and PARP2 on DDR, class switch recombination, PARylation, and different affinities for DSBs [8, 59, 103, 104].

Interestingly, we observed that PARP suppression only increased MMEJ levels when the DSB was induced by ISceI, not Cas9. This differential MMEJ response could be attributed to several reasons. For instance, it could be due to differential enzyme recruitment kinetics and the efficiency of DSB induction. Indeed, we consistently observed higher overall MMEJ levels following Cas9 break induction compared to ISceI (data not shown). Similarly, the mechanisms regulating endonuclease removal from break sites likely differ and could lead to differential effects. It is possible that PARPi-treated Cas9 breaks lead to mild increases in MMEJ levels that are not detected by our system. Conversely, it is likely that the difference between ISceI and Cas9 lies in the type of break and, therefore, could be due to differential DDR responses at blunt vs overhang DSBs. KU70/KU80 heterodimers can strongly bind various DNA structures, including blunt ends, 5’ and 3’ overhangs, and DNA hairpins [105–107]. However, minimal NHEJ factors are needed at blunt DSBs, while more complex breaks require nucleases, polymerases, and accessory factors [108–110].

Notably, NHEJ accessory factor, APLF, interacts with PAR [109, 111, 112]. The increase following PARPi treatment from ISceI breaks may be due to effects on repair factors that process and recognize the 3’ overhangs. Indeed, PARP1 has been implicated in regulating resection at DSBs, though existing data remain contradictory. PARP1 has been shown to promote resection for MMEJ repair in amplicon sequencing experiments following the repair of a Cas9-induced cut [86]. In contrast, others have shown that PARP1 antagonizes or limits resection at breaks [84, 85]. Concordant with all this data, PARP1 can PARylate and/or interact with many proteins known to either promote or prevent resection, including BRCA1, MRE11, KU80, and DNAPKcs, all of which can change the balance in DSB repair pathway determination [6, 85, 113, 114]. Consistent with PARP’s multi-faceted roles in DDR, we observed that PARPi increased MMEJ at ISceI breaks by inhibiting both NHEJ and HR. Together, we predict that PARP inhibition leads to elevated MMEJ activity by dysregulating DDR pathway choice, which can be further exploited via MMEJ inhibition.

Finally, our findings strengthen the rationale for a combined therapeutic approach using PARPi and Polθi in HRD cancers. If PARP1 was required for MMEJ, then further inhibition of MMEJ should be futile. Our findings show that, on the contrary, direct inhibition of MMEJ is likely to work synergistically with PARPi. Moreover, translational data show that Polθ may contribute to PARPi resistance in HRD cancer cells. Indeed, some *BRCA* reversion mutations in diagnosed HRD tumors contain an MMEJ repair signature [28, 29]. We observed increased MMEJ levels of ISceI-induced DSBs following clinically used PARPi drugs in both HR-proficient and *BRCA2*-mutated cells (**Figure 1**). If PARPi inadvertently leads to an upregulation of Polθ-mediated MMEJ for certain types of breaks, which can, in turn, lead to PARPi resistance, then blocking Polθ activity could also act in preventing resistance and thereby lengthening progression-free survival in patients taking PARPi.

## AUTHOR CONTRIBUTIONS

RO, BG and NA designed the study. RO, ET, SW and NA performed experiments. RO, ET, BG, NA analyzed experiments. TD performed bioinformatics analyses. BG and NA supervised the research and acquired funding. RO, BG, NA wrote the manuscript. All authors read and revised the manuscript.

## ACKNOWLEDGEMENTS

We acknowledge philanthropic contributions from Thomas and Kay L. Dunton Endowed Chair in Ovarian Cancer Research. We acknowledge the Light Microscopy Core Facility, Porter Biosciences B047, B049, B051 and B059 at the University of Colorado Boulder (RRID:SCR_018993) for help and advice with microscopy. We thank the Flow Cytometry Core Facility at the University of Colorado Boulder.

This work was supported by the National Institutes of Health [R37CA261987 to BGB with supplement to RO, R01CA266100 to NA, R35GM143108 to NA, P30CA046934 to the University of Colorado Cancer Center]. Funding for open access charge: National Institutes of Health.

## REFERENCES

1. Scully, R., et al., DNA double-strand break repair-pathway choice in somatic mammalian cells. Nat Rev Mol Cell Biol, 2019. 20(11): p. 698–714.

2. Gibson, B.A. and W.L. Kraus, New insights into the molecular and cellular functions of poly(ADP-ribose) and PARPs. Nat Rev Mol Cell Biol, 2012. 13(7): p. 411–24.

3. Langelier, M.F., et al., Structural basis for DNA damage-dependent poly(ADP-ribosyl)ation by human PARP-1. Science, 2012. 336(6082): p. 728–32.

4. Krishnakumar, R. and W.L. Kraus, The PARP side of the nucleus: molecular actions, physiological outcomes, and clinical targets. Mol Cell, 2010. 39(1): p. 8–24.

5. Mortusewicz, O., et al., Feedback-regulated poly(ADP-ribosyl)ation by PARP-1 is required for rapid response to DNA damage in living cells. Nucleic Acids Res, 2007. 35(22): p. 7665–75.

6. Haince, J.F., et al., PARP1-dependent kinetics of recruitment of MRE11 and NBS1 proteins to multiple DNA damage sites. J Biol Chem, 2008. 283(2): p. 1197–208.

7. Schreiber, V., et al., Poly(ADP-ribose): novel functions for an old molecule. Nat Rev Mol Cell Biol, 2006. 7(7): p. 517–28.

8. Matta, E., et al., Insight into DNA substrate specificity of PARP1-catalysed DNA poly(ADP-ribosyl)ation. Sci Rep, 2020. 10(1): p. 3699.

9. Beck, C., et al., Poly(ADP-ribose) polymerases in double-strand break repair: focus on PARP1, PARP2 and PARP3. Exp Cell Res, 2014. 329(1): p. 18–25.

10. Farmer, H., et al., Targeting the DNA repair defect in BRCA mutant cells as a therapeutic strategy. Nature, 2005. 434(7035): p. 917-21.

11. Bryant, H.E., et al., Specific killing of BRCA2-deficient tumours with inhibitors of poly(ADP-ribose) polymerase. Nature, 2005. 434(7035): p. 913–7.

12. Li, X. and L. Zou, BRCAness, DNA gaps, and gain and loss of PARP inhibitor-induced synthetic lethality. J Clin Invest, 2024. 134(14).

13. Bhamidipati, D., et al., PARP inhibitors: enhancing efficacy through rational combinations. Br J Cancer, 2023. 129(6): p. 904–916.

14. Chaudhuri, A.R., et al., Erratum: Replication fork stability confers chemoresistance in BRCA-deficient cells. Nature, 2016. 539(7629): p. 456.

15. Cong, K., et al., Replication gaps are a key determinant of PARP inhibitor synthetic lethality with BRCA deficiency. Mol Cell, 2021. 81(15): p. 3227.

16. Murai, J., et al., Trapping of PARP1 and PARP2 by Clinical PARP Inhibitors. Cancer Res, 2012. 72(21): p. 5588–99.

17. Panzarino, N.J., et al., Replication Gaps Underlie BRCA Deficiency and Therapy Response. Cancer Res, 2021. 81(5): p. 1388–1397.

18. Schlacher, K., et al., Double-strand break repair-independent role for BRCA2 in blocking stalled replication fork degradation by MRE11. Cell, 2011. 145(4): p. 529–42.

19. Schlacher, K., H. Wu, and M. Jasin, A distinct replication fork protection pathway connects Fanconi anemia tumor suppressors to RAD51-BRCA1/2. Cancer Cell, 2012. 22(1): p. 106–16.

20. Simoneau, A., R. Xiong, and L. Zou, The trans cell cycle effects of PARP inhibitors underlie their selectivity toward BRCA1/2-deficient cells. Genes Dev, 2021. 35(17-18): p. 1271–1289.

21. Vaitsiankova, A., et al., PARP inhibition impedes the maturation of nascent DNA strands during DNA replication. Nat Struct Mol Biol, 2022. 29(4): p. 329–338.

22. Audeh, M.W., et al., Oral poly(ADP-ribose) polymerase inhibitor olaparib in patients with BRCA1 or BRCA2 mutations and recurrent ovarian cancer: a proof-of-concept trial. Lancet, 2010. 376(9737): p. 245–51.

23. Fong, P.C., et al., Poly(ADP)-ribose polymerase inhibition: frequent durable responses in BRCA carrier ovarian cancer correlating with platinum-free interval. J Clin Oncol, 2010. 28(15): p. 2512–9.

24. Li, H., et al., PARP inhibitor resistance: the underlying mechanisms and clinical implications. Mol Cancer, 2020. 19(1): p. 107.

25. Yazinski, S.A., et al., ATR inhibition disrupts rewired homologous recombination and fork protection pathways in PARP inhibitor-resistant BRCA-deficient cancer cells. Genes Dev, 2017. 31(3): p. 318–332.

26. Pettitt, S.J., et al., Genome-wide and high-density CRISPR-Cas9 screens identify point mutations in PARP1 causing PARP inhibitor resistance. Nat Commun, 2018. 9(1): p. 1849.

27. Christie, E.L., et al., Multiple ABCB1 transcriptional fusions in drug resistant high-grade serous ovarian and breast cancer. Nat Commun, 2019. 10(1): p. 1295.

28. Tobalina, L., et al., A meta-analysis of reversion mutations in BRCA genes identifies signatures of DNA end-joining repair mechanisms driving therapy resistance. Ann Oncol, 2021. 32(1): p. 103–112.

29. Pettitt, S.J., et al., Clinical BRCA1/2 Reversion Analysis Identifies Hotspot Mutations and Predicted Neoantigens Associated with Therapy Resistance. Cancer Discov, 2020. 10(10): p. 1475–1488.

30. Watson, Z.L., et al., Histone methyltransferases EHMT1 and EHMT2 (GLP/G9A) maintain PARP inhibitor resistance in high-grade serous ovarian carcinoma. Clinical Epigenetics, 2019. 11(1).

31. Nguyen, L.L., et al., EHMT1/2 inhibition promotes regression of therapy-resistant ovarian cancer tumors in a CD8 T cell-dependent manner. Mol Cancer Res, 2024.

32. Nguyen, L.L., et al., Combining EHMT and PARP Inhibition: A Strategy to Diminish Therapy-Resistant Ovarian Cancer Tumor Growth while Stimulating Immune Activation. Mol Cancer Ther, 2024: p. OF1–OF16.

33. Bouwman, P., et al., 53BP1 loss rescues BRCA1 deficiency and is associated with triple-negative and BRCA-mutated breast cancers. Nat Struct Mol Biol, 2010. 17(6): p. 688–95.

34. Bunting, S.F., et al., 53BP1 inhibits homologous recombination in Brca1-deficient cells by blocking resection of DNA breaks. Cell, 2010. 141(2): p. 243–54.

35. Zhou, J., et al., A first-in-class Polymerase Theta Inhibitor selectively targets Homologous-Recombination-Deficient Tumors. Nat Cancer, 2021. 2(6): p. 598–610.

36. Mateos-Gomez, P.A., et al., Mammalian polymerase theta promotes alternative NHEJ and suppresses recombination. Nature, 2015. 518(7538): p. 254–7.

37. Ceccaldi, R., et al., Homologous-recombination-deficient tumours are dependent on Polθ-mediated repair. Nature, 2015. 518(7538): p. 258–262.

38. Zatreanu, D., et al., Poltheta inhibitors elicit BRCA-gene synthetic lethality and target PARP inhibitor resistance. Nat Commun, 2021. 12(1): p. 3636.

39. Schrempf, A., J. Slyskova, and J.I. Loizou, Targeting the DNA Repair Enzyme Polymerase theta in Cancer Therapy. Trends Cancer, 2021. 7(2): p. 98–111.

40. Truong, L.N., et al., Microhomology-mediated End Joining and Homologous Recombination share the initial end resection step to repair DNA double-strand breaks in mammalian cells. Proc Natl Acad Sci U S A, 2013. 110(19): p. 7720–5.

41. Bennardo, N., et al., Alternative-NHEJ is a mechanistically distinct pathway of mammalian chromosome break repair. PLoS Genet, 2008. 4(6): p. e1000110.

42. Newman, J.A., et al., Structure of the Helicase Domain of DNA Polymerase Theta Reveals a Possible Role in the Microhomology-Mediated End-Joining Pathway. Structure, 2015. 23(12): p. 2319–2330.

43. Fijen, C., et al., Sequential requirements for distinct Poltheta domains during theta-mediated end joining. Mol Cell, 2024. 84(8): p. 1460–1474 e6.

44. Fleury, H., et al., The APE2 nuclease is essential for DNA double-strand break repair by microhomology-mediated end joining. Mol Cell, 2023. 83(9): p. 1429–1445 e8.

45. Bai, W., et al., The 3’-flap endonuclease XPF-ERCC1 promotes alternative end joining and chromosomal translocation during B cell class switching. Cell Rep, 2021. 36(13): p. 109756.

46. Kent, T., et al., Mechanism of microhomology-mediated end-joining promoted by human DNA polymerase theta. Nat Struct Mol Biol, 2015. 22(3): p. 230–7.

47. Zahn, K.E., et al., Human DNA polymerase theta grasps the primer terminus to mediate DNA repair. Nat Struct Mol Biol, 2015. 22(4): p. 304–11.

48. Crespan, E., et al., Microhomology-mediated DNA strand annealing and elongation by human DNA polymerases lambda and beta on normal and repetitive DNA sequences. Nucleic Acids Res, 2012. 40(12): p. 5577–90.

49. Chandramouly, G., et al., Pollambda promotes microhomology-mediated end-joining. Nat Struct Mol Biol, 2023. 30(1): p. 107–114.

50. Stroik, S., et al., Stepwise requirements for polymerases delta and theta in theta-mediated end joining. Nature, 2023. 623(7988): p. 836-841.

51. Wang, H., et al., DNA ligase III as a candidate component of backup pathways of nonhomologous end joining. Cancer Res, 2005. 65(10): p. 4020–30.

52. Simsek, D., et al., DNA ligase III promotes alternative nonhomologous end-joining during chromosomal translocation formation. PLoS Genet, 2011. 7(6): p. e1002080.

53. Lu, G., et al., Ligase I and ligase III mediate the DNA double-strand break ligation in alternative end-joining. Proc Natl Acad Sci U S A, 2016. 113(5): p. 1256–60.

54. Paul, K., et al., DNA ligases I and III cooperate in alternative non-homologous end-joining in vertebrates. PLoS One, 2013. 8(3): p. e59505.

55. Audebert, M., B. Salles, and P. Calsou, Involvement of poly(ADP-ribose) polymerase-1 and XRCC1/DNA ligase III in an alternative route for DNA double-strand breaks rejoining. J Biol Chem, 2004. 279(53): p. 55117–26.

56. Wang, M., et al., PARP-1 and Ku compete for repair of DNA double strand breaks by distinct NHEJ pathways. Nucleic Acids Res, 2006. 34(21): p. 6170–82.

57. Jia, Q., et al., Poly(ADP-ribose)polymerases are involved in microhomology mediated back-up non-homologous end joining in Arabidopsis thaliana. Plant Mol Biol, 2013. 82(4-5): p. 339–51.

58. Sharma, S., et al., Homology and enzymatic requirements of microhomology-dependent alternative end joining. Cell Death Dis, 2015. 6(3): p. e1697.

59. Robert, I., F. Dantzer, and B. Reina-San-Martin, Parp1 facilitates alternative NHEJ, whereas Parp2 suppresses IgH/c-myc translocations during immunoglobulin class switch recombination. J Exp Med, 2009. 206(5): p. 1047–56.

60. Soni, A., et al., Requirement for Parp-1 and DNA ligases 1 or 3 but not of Xrcc1 in chromosomal translocation formation by backup end joining. Nucleic Acids Res, 2014. 42(10): p. 6380–92.

61. Sfeir, A. and T. de Lange, Removal of shelterin reveals the telomere end-protection problem. Science, 2012. 336(6081): p. 593–7.

62. Kais, Z., et al., FANCD2 Maintains Fork Stability in BRCA1/2-Deficient Tumors and Promotes Alternative End-Joining DNA Repair. Cell Rep, 2016. 15(11): p. 2488–99.

63. Yun, M.H. and K. Hiom, CtIP-BRCA1 modulates the choice of DNA double-strand-break repair pathway throughout the cell cycle. Nature, 2009. 459(7245): p. 460–3.

64. Mateos-Gomez, P.A., et al., The helicase domain of Poltheta counteracts RPA to promote alt-NHEJ. Nat Struct Mol Biol, 2017. 24(12): p. 1116-1123.

65. Chen, L., et al., Cell cycle-dependent complex formation of BRCA1.CtIP.MRN is important for DNA double-strand break repair. J Biol Chem, 2008. 283(12): p. 7713–20.

66. Ahrabi, S., et al., A role for human homologous recombination factors in suppressing microhomology-mediated end joining. Nucleic Acids Res, 2016. 44(12): p. 5743–57.

67. Wang, H., et al., PLK1 targets CtIP to promote microhomology-mediated end joining. Nucleic Acids Res, 2018. 46(20): p. 10724–10739.

68. Brambati, A., et al., RHINO directs MMEJ to repair DNA breaks in mitosis. Science, 2023. 381(6658): p. 653–660.

69. Gelot, C., et al., Poltheta is phosphorylated by PLK1 to repair double-strand breaks in mitosis. Nature, 2023. 621(7978): p. 415–422.

70. Lee, D.H., et al., Dephosphorylation enables the recruitment of 53BP1 to double-strand DNA breaks. Mol Cell, 2014. 54(3): p. 512–25.

71. Ceccaldi, R., B. Rondinelli, and A.D. D’Andrea, Repair Pathway Choices and Consequences at the Double-Strand Break. Trends Cell Biol, 2016. 26(1): p. 52–64.

72. Blackford, A.N. and M. Stucki, How Cells Respond to DNA Breaks in Mitosis. Trends Biochem Sci, 2020. 45(4): p. 321–331.

73. Terasawa, M., A. Shinohara, and M. Shinohara, Canonical non-homologous end joining in mitosis induces genome instability and is suppressed by M-phase-specific phosphorylation of XRCC4. PLoS Genet, 2014. 10(8): p. e1004563.

74. Llorens-Agost, M., et al., POLtheta-mediated end joining is restricted by RAD52 and BRCA2 until the onset of mitosis. Nat Cell Biol, 2021. 23(10): p. 1095–1104.

75. Taylor, W.R., FACS-based detection of phosphorylated histone H3 for the quantitation of mitotic cells. Methods Mol Biol, 2004. 281: p. 293–9.

76. Ledermann, J.A., PARP inhibitors in ovarian cancer. Ann Oncol, 2016. 27 Suppl 1: p. i40–i44.

77. Del Conte, G., et al., Phase I study of olaparib in combination with liposomal doxorubicin in patients with advanced solid tumours. Br J Cancer, 2014. 111(4): p. 651–9.

78. Sfeir, A., M. Tijsterman, and M. McVey, Microhomology-Mediated End-Joining Chronicles: Tracing the Evolutionary Footprints of Genome Protection. Annu Rev Cell Dev Biol, 2024. 40(1): p. 195–218.

79. Jelinic, P. and D.A. Levine, New insights into PARP inhibitors’ effect on cell cycle and homology-directed DNA damage repair. Mol Cancer Ther, 2014. 13(6): p. 1645–54.

80. Chan, S.H., A.M. Yu, and M. McVey, Dual roles for DNA polymerase theta in alternative end-joining repair of double-strand breaks in Drosophila. PLoS Genet, 2010. 6(7): p. e1001005.

81. Yousefzadeh, M.J., et al., Mechanism of suppression of chromosomal instability by DNA polymerase POLQ. PLoS Genet, 2014. 10(10): p. e1004654.

82. Rudolph, J., K. Jung, and K. Luger, Inhibitors of PARP: Number crunching and structure gazing. Proc Natl Acad Sci U S A, 2022. 119(11): p. e2121979119.

83. Thorsell, A.G., et al., Structural Basis for Potency and Promiscuity in Poly(ADP-ribose) Polymerase (PARP) and Tankyrase Inhibitors. J Med Chem, 2017. 60(4): p. 1262–1271.

84. Caron, M.C., et al., Poly(ADP-ribose) polymerase-1 antagonizes DNA resection at double-strand breaks. Nat Commun, 2019. 10(1): p. 2954.

85. Lodovichi, S., et al., PARylation of BRCA1 limits DNA break resection through BRCA2 and EXO1. Cell Rep, 2023. 42(2): p. 112060.

86. Luedeman, M.E., et al., Poly(ADP) ribose polymerase promotes DNA polymerase theta-mediated end joining by activation of end resection. Nat Commun, 2022. 13(1): p. 4547.

87. Lei, T., et al., Multifaceted regulation and functions of 53BP1 in NHEJ-mediated DSB repair (Review). Int J Mol Med, 2022. 50(1).

88. Hoa, N.N., et al., Relative contribution of four nucleases, CtIP, Dna2, Exo1 and Mre11, to the initial step of DNA double-strand break repair by homologous recombination in both the chicken DT40 and human TK6 cell lines. Genes Cells, 2015. 20(12): p. 1059–76.

89. Nimonkar, A.V., et al., BLM-DNA2-RPA-MRN and EXO1-BLM-RPA-MRN constitute two DNA end resection machineries for human DNA break repair. Genes Dev, 2011. 25(4): p. 350–62.

90. Zhao, F., et al., DNA end resection and its role in DNA replication and DSB repair choice in mammalian cells. Exp Mol Med, 2020. 52(10): p. 1705–1714.

91. Sturzenegger, A., et al., DNA2 cooperates with the WRN and BLM RecQ helicases to mediate long-range DNA end resection in human cells. J Biol Chem, 2014. 289(39): p. 27314–27326.

92. Symington, L.S., Mechanism and regulation of DNA end resection in eukaryotes. Crit Rev Biochem Mol Biol, 2016. 51(3): p. 195–212.

93. Hayashi, M.T., et al., A telomere-dependent DNA damage checkpoint induced by prolonged mitotic arrest. Nat Struct Mol Biol, 2012. 19(4): p. 387–94.

94. Kwon, M., M.L. Leibowitz, and J.H. Lee, Small but mighty: the causes and consequences of micronucleus rupture. Exp Mol Med, 2020. 52(11): p. 1777–1786.

95. Wang, J., C.A. Sadeghi, and R.L. Frock, DNA-PKcs suppresses illegitimate chromosome rearrangements. Nucleic Acids Res, 2024. 52(9): p. 5048–5066.

96. Liang, Z., et al., Ku70 suppresses alternative end joining in G1-arrested progenitor B cells. Proc Natl Acad Sci U S A, 2021. 118(21).

97. Vergara, X., et al., Widespread chromatin context-dependencies of DNA double-strand break repair proteins. Nat Commun, 2024. 15(1): p. 5334.

98. Yu, W., et al., Repair of G1 induced DNA double-strand breaks in S-G2/M by alternative NHEJ. Nat Commun, 2020. 11(1): p. 5239.

99. Schaub, J.M., M.M. Soniat, and I.J. Finkelstein, Polymerase theta-helicase promotes end joining by stripping single-stranded DNA-binding proteins and bridging DNA ends. Nucleic Acids Res, 2022. 50(7): p. 3911–3921.

100. Mansour, W.Y., et al., The absence of Ku but not defects in classical non-homologous end-joining is required to trigger PARP1-dependent end-joining. DNA Repair (Amst), 2013. 12(12): p. 1134–42.

101. El-Khamisy, S.F., et al., A requirement for PARP-1 for the assembly or stability of XRCC1 nuclear foci at sites of oxidative DNA damage. Nucleic Acids Res, 2003. 31(19): p. 5526–33.

102. Vekariya, U., et al., PARG is essential for Poltheta-mediated DNA end-joining by removing repressive poly-ADP-ribose marks. Nat Commun, 2024. 15(1): p. 5822.

103. Langelier, M.F., A.A. Riccio, and J.M. Pascal, PARP-2 and PARP-3 are selectively activated by 5’ phosphorylated DNA breaks through an allosteric regulatory mechanism shared with PARP-1. Nucleic Acids Res, 2014. 42(12): p. 7762–75.

104. Obaji, E., T. Haikarainen, and L. Lehtio, Structural basis for DNA break recognition by ARTD2/PARP2. Nucleic Acids Res, 2018. 46(22): p. 12154–12165.

105. Mimori, T. and J.A. Hardin, Mechanism of interaction between Ku protein and DNA. J Biol Chem, 1986. 261(22): p. 10375–9.

106. Paillard, S. and F. Strauss, Analysis of the mechanism of interaction of simian Ku protein with DNA. Nucleic Acids Res, 1991. 19(20): p. 5619–24.

107. Doherty, A.J. and S.P. Jackson, DNA repair: how Ku makes ends meet. Curr Biol, 2001. 11(22): p. R920–4.

108. Ma, Y., K. Schwarz, and M.R. Lieber, The Artemis:DNA-PKcs endonuclease cleaves DNA loops, flaps, and gaps. DNA Repair (Amst), 2005. 4(7): p. 845–51.

109. Li, S., et al., Polynucleotide kinase and aprataxin-like forkhead-associated protein (PALF) acts as both a single-stranded DNA endonuclease and a single-stranded DNA 3’ exonuclease and can participate in DNA end joining in a biochemical system. J Biol Chem, 2011. 286(42): p. 36368–77.

110. Chang, H.H.Y., et al., Different DNA End Configurations Dictate Which NHEJ Components Are Most Important for Joining Efficiency. J Biol Chem, 2016. 291(47): p. 24377–24389.

111. Eustermann, S., et al., Solution structures of the two PBZ domains from human APLF and their interaction with poly(ADP-ribose). Nat Struct Mol Biol, 2010. 17(2): p. 241–3.

112. Rulten, S.L., et al., PARP-3 and APLF function together to accelerate nonhomologous end-joining. Mol Cell, 2011. 41(1): p. 33–45.

113. Han, Y., et al., DNA-PKcs PARylation regulates DNA-PK kinase activity in the DNA damage response. Mol Med Rep, 2019. 20(4): p. 3609–3616.

114. Sajish, M., et al., Trp-tRNA synthetase bridges DNA-PKcs to PARP-1 to link IFN-gamma and p53 signaling. Nat Chem Biol, 2012. 8(6): p. 547–54.

