## Supplemental Figures for "PARP1 and PARP2 are dispensable for DNA repair by microhomology-mediated end-joining during mitosis"

† Joint Authors

* To whom correspondence should be addressed:

* Correspondence may also be addressed to:

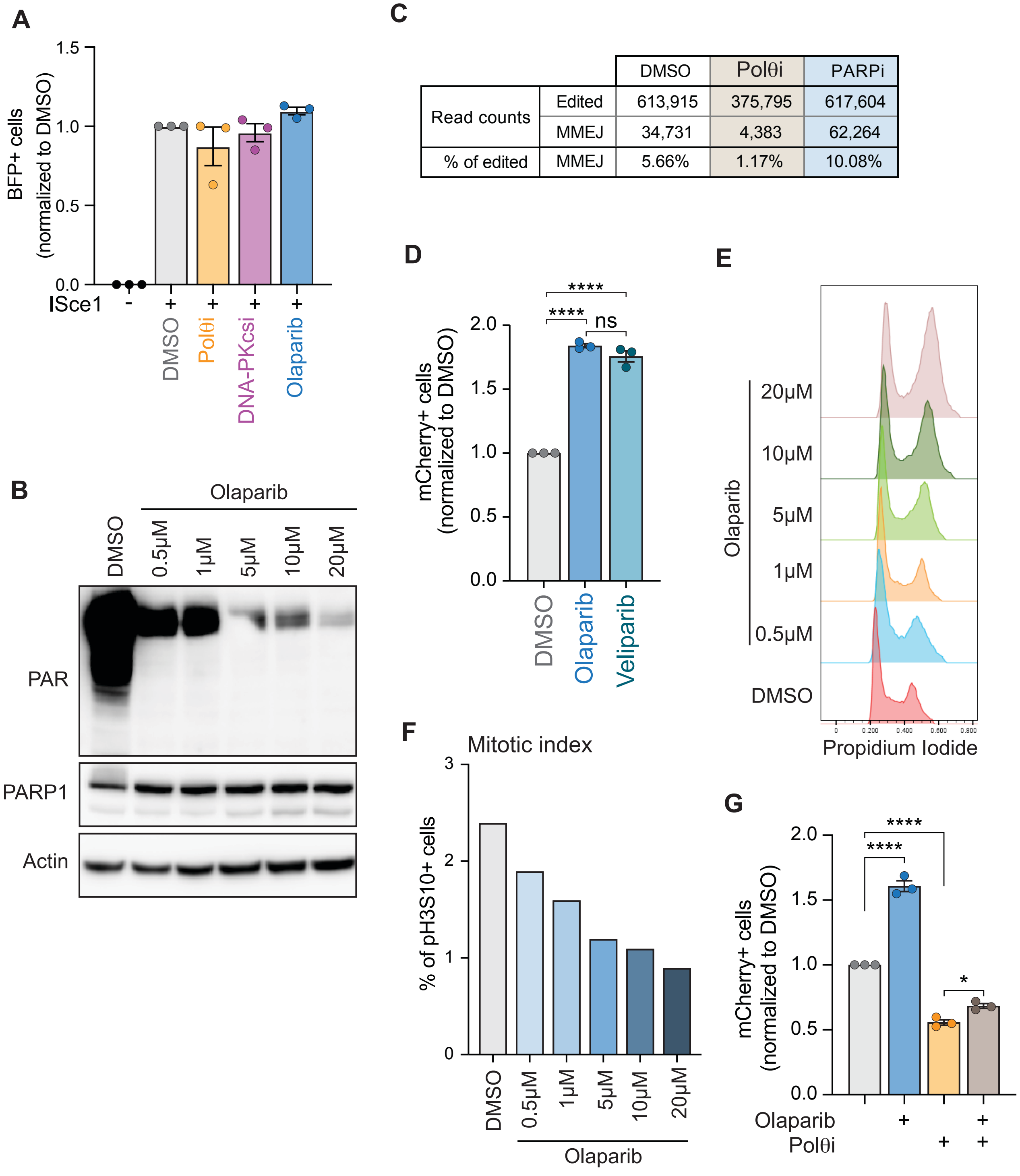


| | | DMSO | Polθi | PARPi |
|---|---|---|---|---|
| Read counts | Edited | 613,915 | 375,795 | 617,604 |
| | MMEJ | 34,731 | 4,383 | 62,264 |
| % of edited | MMEJ | 5.66% | 1.17% | 10.08% |

**Ortega et al., Supplementary Figure 1**

**Supplementary Figure 1.**

(**A**) BFP+ (IScel+) quantification from Figure 1C in HT1080 cells. Values are normalized to DMSO. (**B**) Immunoblot of PARP1, Actin, and PAR in dose escalated olaparib HT1080 cells. (**C**) Table of quantification of amplicon sequencing from data in Figure 1C to validate flow cytometry results. Data represent percent of edited sequences, one experiment. (**D**) MMEJ quantification using reporter and timeline from Figure 1A-B in HT1080 cells following olaparib (5 μm) or veliparib (5 μm) treatment. Values are normalized to DMSO. (**E**) Flow cytometry cell cycle analysis of dose escalated olaparib cells using propidium iodide staining. (**F**) Flow cytometry mitotic index using pH3S10+ cells following dose escalated olaparib. (**G**) MMEJ quantification using reporter and timeline from Figure 1A-B in HT1080 cells following olaparib (5 μm), Polθi (ART558, 10 μm) or combination. Values are normalized to DMSO. Statistical analyses for (**D and G**): Data represent three independent experiments, each the average of three technical replicates. Data are mean ± SEM. Statistical test, one way ANOVA with multiple comparison correction. ns: non-significant, *p<0.05, ****p<0.0001.

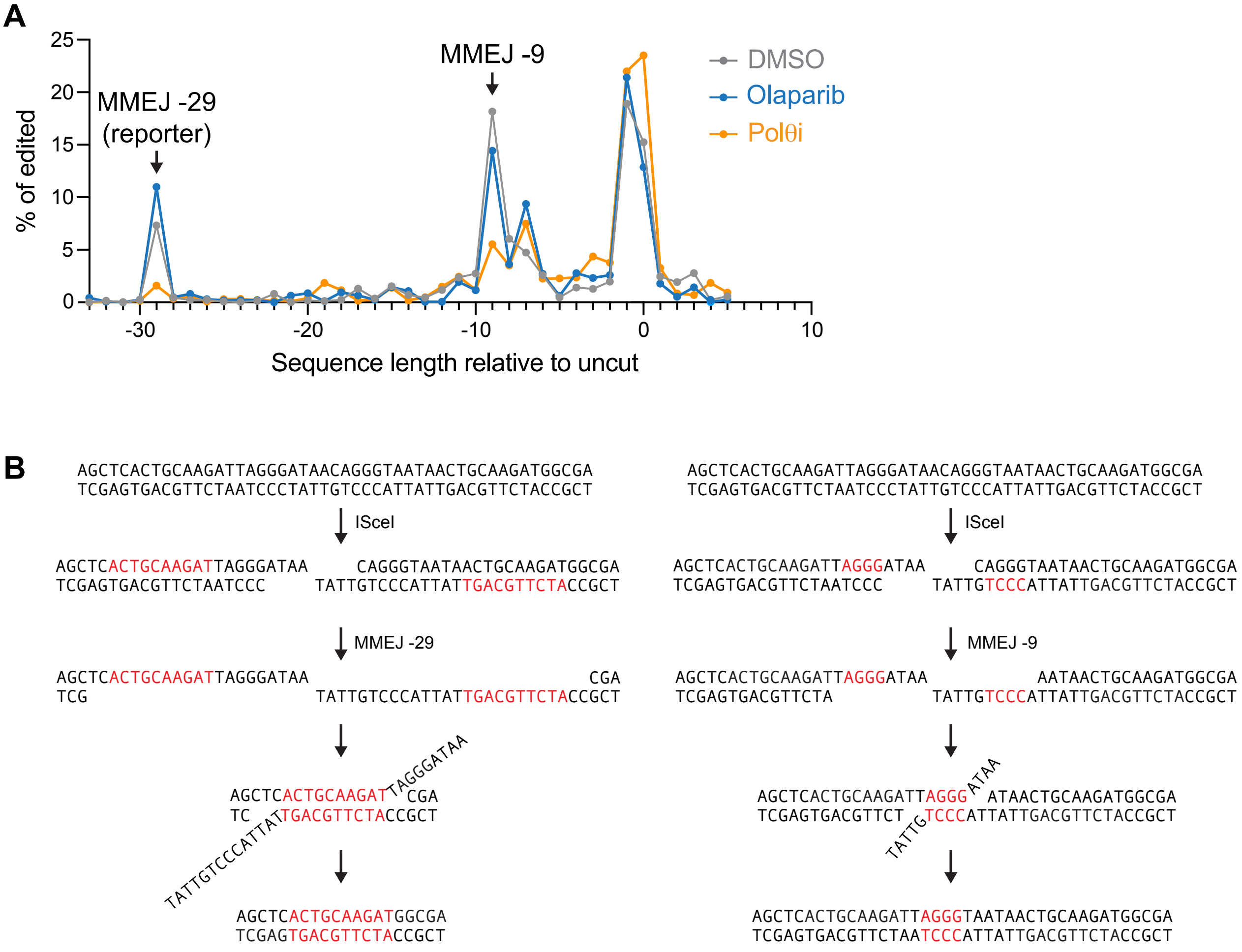


**Ortega et al., Supplementary Figure 2**

**Supplementary Figure 2.**

(**A**) Amplicon sequencing from MMEJ reporter and timeline from Figure 1A-B in HT1080 cells following DMSO, Polθi (ART558, 10 μm), or olaparib (5 μm) treatment. Graph shows all edited products sequence lengths relative to unedited. mCherry+ product corresponds to the -29 nucleotides deletion. (**B**) Schematic of sequence and repair for the -29 (left) and -9 (right) deletions. Red depicts microhomologies.

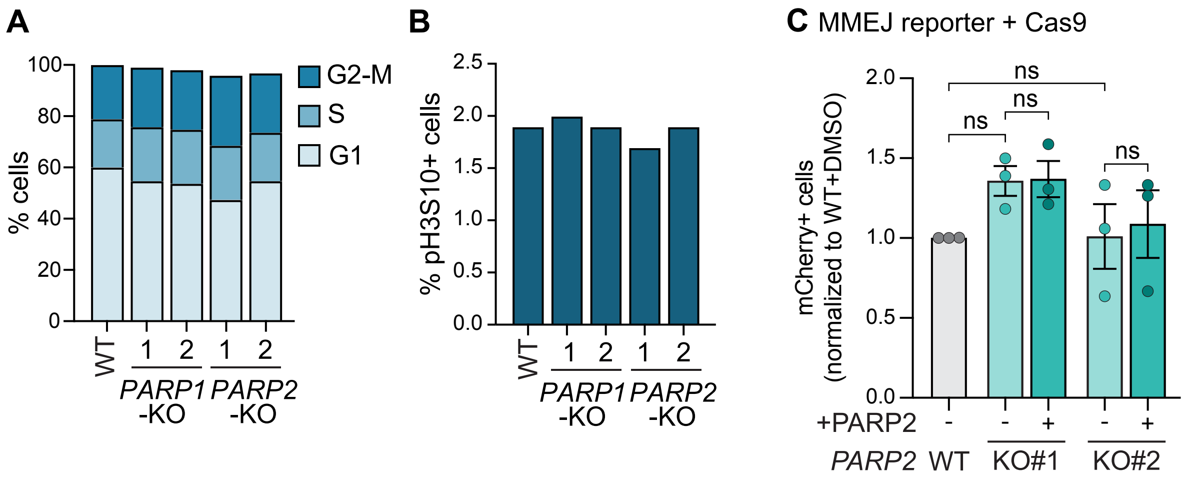


**Ortega et al., Supplementary Figure 3**

**Supplementary Figure 3.**

(**A**) Flow cytometry cell cycle analysis using propidium iodide (PI) of cells used in (C), Figure 2C-D, and Figure 5C-E. (**B**) Flow cytometry mitotic index using pH3S10+ of cells used in (C), Figure 2C-D, and Figure 5C-E. (**C**) MMEJ quantification using the reporter with Cas9 cut (from Figure 4A-B) in HT1080 cells and isogenic *PARP2*-KO cells with or without complemented *PARP2*. Values are normalized to wild-type DMSO. Statistical analyses for (**C**): Data represent three independent experiments. Data are mean ± SEM. Statistical test, one way ANOVA with multiple comparison correction. ns: non-significant.
